# The transcriptional regulatory circuit as a driver and therapeutic target in CML blast crisis

**DOI:** 10.1101/2025.11.02.686161

**Authors:** Liling Jiang, Enzhe Lou, Jinxin Fang, Lizhen Jiang, Guanjie Peng, Jinghong Chen, Bingyuan Liu, Bo Lu, Yi Meng, Haichuan Zhang, Aochu Liu, Qiong Mao, Peilong Lai, Yueyuan Zheng, Jinbao Liu, Xianping Shi

## Abstract

Patients with blast crisis in chronic myeloid leukemia (CML) exhibit an extremely poor prognosis, with median survival of only a few months and limited therapeutic options. Dysregulated transcriptional regulation plays a pivotal role in driving blast crisis progression. In this study, we performed histone modification and transcriptomic high-throughput sequencing on bone marrow cells derived from chronic and blast-phase CML patients, identifying a CML blast-specific transcriptional regulatory circuit comprising MEF2C, MYB, MEIS1, and ZEB2. These transcription factors (TFs) form protein complexes that co-bind to shared enhancers or promoters, reciprocally enhancing each other’s transcriptional activity and that of their downstream targets. Mechanistically, this circuit synergistically reprograms chromatin accessibility and transcriptional landscapes in blast cells, parallel activation of purine metabolism and Rho GTPase signaling pathways which exhibit functional crosstalk. Functional validation demonstrated that silencing components of this circuit or its downstream core targets significantly suppressed CML blast progression *in vitro* and *in vivo*. Notably, mebendazole (MBZ), a clinically approved inhibitor targeting MYB protein stability, recapitulated these inhibitory effects. Our findings establish the MEF2C-MYB-MEIS1-ZEB2 transcriptional circuit as a central driver of CML blast crisis and position MBZ as a high-potential translational strategy for targeting this lethal phase of the disease.

**Key Points:**

1. MEF2C, MYB, MEIS1, and ZEB2 form the CML blast-crisis transcriptional circuit, synergistically reprogramming chromatin and transcription.
2. Mebendazole inhibits CML blast progression by impairing the circuit, offering a rapid translational strategy for this lethal phase.

## Introduction

Chronic myeloid leukemia (CML) has seen remarkable therapeutic advances with the introduction of tyrosine kinase inhibitors (TKIs); nevertheless, a subset of patients still progresses to blast-phase CML (BP-CML). A hallmark of BP-CML is impaired differentiation of primitive hematopoietic progenitors, with bone marrow or peripheral blood blasts <10% in chronic phase and >20% in blast phase^1^. Current therapeutic options for BP-CML remain highly limited, with a median overall survival of only 2-3 months^2^. The pathogenesis of blast crisis progression in CML is a complex, heterogeneous process driven by multiple interconnected mechanisms. Prior investigations have largely centered on genomic alterations arising from BCR-ABL amplification, mutations in hematopoietic lineage transcription factors, and epigenetic dysregulation, all of which contribute to defective differentiation or uncontrolled proliferation^3-7^. These studies have primarily focused on elucidating single-gene roles, yet they are hampered by significant heterogeneity and challenges in translating candidate targets to clinical use. Growing evidence highlights that super-enhancer-driven transcriptional regulatory networks, which orchestrate cohorts of transcription factors, play pivotal roles in establishing cellular identity and propelling disease onset/progression^8, 9^. However, whether a core transcriptional regulatory network of transcription factors governs the progression of CML to blast crisis remains unclear.

Among all transcription factors (TFs), nearly 50% are ubiquitously expressed across eukaryotic cell types, whereas a restricted subset dictates cellular identity by orchestrating specific gene expression programs^10^. This latter group often driven by super enhancers (SEs), which serve as pivotal controllers of cellular gene expression^8^. Emerging evidence demonstrates that these SE-driving TFs typically form interconnected regulatory networks, with interactions spanning direct transcriptional regulation and protein-protein interactions^11, 12^. These networks act hierarchically to drive the expression of cell-identity-defining genes, playing critical roles in maintaining cell-type-specific transcriptional programs and biological functions^11,13-15^.

Preclinical and clinical studies have established that targeting these transcriptional regulatory networks represents a potent therapeutic strategy for cancer^16-18^. Oncogenes governed by these circuits exhibit heightened dependence on sustained transcriptional activation, rendering them exquisitely sensitive to interventions that disrupt transcriptional machinery. This vulnerability underscores the significant promise of developing novel targeted therapies that specifically perturb these networks^13, 19^. Thus, from the perspective of core transcriptional regulation, identifying novel tumor driver genes and targeting their regulatory networks is poised to emerge as a powerful approach in cancer therapeutics.

Our study employed H3K27ac ChIP-seq and RNA-seq on CP- and BP-CMLs to characterize blast phase-specific SE-driven TFs. Four TFs including MEF2C, MYB, MEIS1, ZEB2 were validated to reciprocally regulate each other via enhancer/promoter binding, collectively governing chromatin accessibility and transcriptional programs in BP cells. These TFs were validated to promote BP progression both *in vitro* and *in vivo*. Treatment with Mebendazol, suppressed the expression of other components within this regulatory circuit. Notably, Mebendazol not only inhibited the growth of BP cells but also mitigated disease progression and extended survival in BP mouse models.

## Methods and Materials

For full details of methods, please refer to the supplemental Material.

### Cell culture

Human CML cell lines (K562, KU812, LAMA84, KBM5) and human embryonic kidney 293T cells were used in this study. 293T cells were cultured in gibco’s medium (DMEM) containing 10% fetal bovine serum protein. K562, KU812 and LAMA84 were cultured in gibco’s medium (1640) containing 10% fetal bovine serum protein. KBM5 was grown in gibco’s medium (IMDM) containing 10% fetal bovine serum protein.

### Isolation of mononuclear cells

Bone marrow cells were collected from patients at The Seventh Affiliated Hospital of Sun Yat-sen University with institutional review board (IRB) approval (approve number: KY-2025-010-01). Mononuclear cells were isolated from the buffy coat layer following density gradient centrifugation. Subsequently, red blood cells were lysed using pre-cooled red blood cell (RBC) lysis buffer (155 mM NH_4_Cl, 10 mM KHCO_3_, 0.1 mM EDTA) for 5 minutes at room temperature. Cell pellets were collected via centrifugation at 1000 rpm for 5 minutes, washed twice with PBS, and re-pelleted under identical centrifugation conditions. The detailed case information collected in this study is provided in Supplementary table 1.

### Super-enhancer analysis

BP-specific peaks were analyzed using DiffBind v3.4.11, applying a cutoff of |log2FC| > 0.584 and FDR < 0.05. Following this, SE were identified with ROSE, which merged enhancers within a 12 kb distance and called SEs based on signal-intensity inflection point (cutoff >1). Transcription factors associated with BP super-enhancers in each sample were then consolidated and selected based on their upregulation in BP tissues.

## Results

### 1. A transcriptional regulation circuit identified in blast CML

To compare the differences in the super-enhancer landscape between CP-CML and BP-CMP, we performed the ChIP-Seq targeting H3K27ac in bone marrow mononuclear cell of CML patients. We screened 6152 BP specific peaks (Fig. 1A), and identified 465 SE associated genes, which including 56 transcription factors (TFs) (Fig. 1B). Among the 56 SE driven TFs, 16 TFs were confirmed to be highly expressed in BP-CML across in house and Branford et al. RNA sequencing datasets^3^(Fig. 1C). Furthermore, we performed a preliminary functional screening of 16 SE-TFs using CRIPRS scores of BP-CML cells from the CCLE DepMap database, identifying 10 SE-TFs with scores less than 0 (Fig. 1D). These TFs are highly expressed in BP-CML patients which verified by two independently RNA-Seq datasets and one microarry dadaset^3, 20^(Fig. S1A). Notably, these TFs also exhibited higher expression in the BP-CML compared with its paired CP (Fig. S1B). The chromatin accessibility at the loci of these TFs is higher in the BP-CML (Fig. S1C). There are included MYB, RUNX1, MEIS1, and HOXA9—previously reported to promote CML blast crisis progression ^21-23^. Since core SE-TFs typically exhibit transcriptional and protein-level interactions that synergistically define cellular identity in loop configurations, we performed transcriptional regulation and protein-protein interaction predictions for the 10 screened SE-TFs. The results suggest that ZEB2, MEIS1, MYB, MEF2C, HOXA9, SOX4, and RUNX1 may composed a regulation circuit (Fig. 1E, 1F).

**Figure 1.**
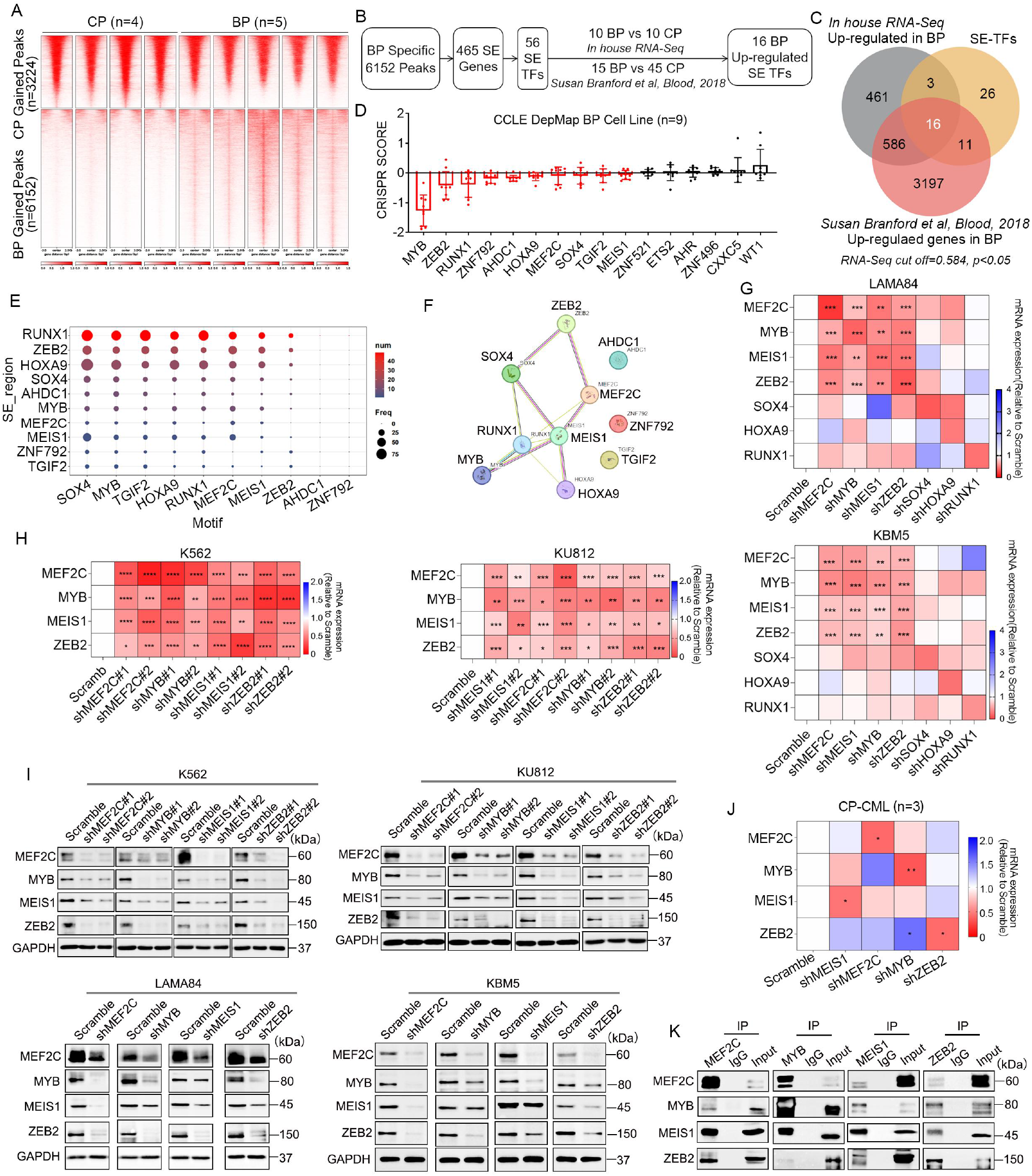
The transcriptional regulation loop are composed of MEF2C, MYB, MEIS1, and ZEB2. A. Differential expression peaks between BP-CML (n=5) and CP-CML (n=4). B. Flowchart of BP-specific SE-TF screening. C. Intersection of SE-TFs commonly overexpressed in BP-CML across three independent datasets. D. CRISPR scores of 16 candidate SE-TFs in BP-CML cell lines (n=9). E. Binding frequency of 10 candidate SE-TFs in reciprocal SE regions. F. Protein-protein reciprocal regulatory interactions among 10 candidate SE-TFs predicted by the STRING database. G,H. In BP-CML cells, individual knockdown of seven candidate SE-TFs was performed to detect their mRNA expression. I. In BP-CML cells, individual knockdown of candidate SE-TFs was performed to detect their protein expression. J. Following SE-TF knockdown in peripheral blood mononuclear cells from CP-CML patients, the mRNA expression of each TF was detected. K. Co-IP assays were performed to detect protein-protein interactions among the four SE-TFs.

To validate the existence of the regulatory circuit, we knocked down the seven candidate SE-TFs individually in BP-CML cell lines and examined their mRNA expression. The results revealed mutual regulatory relationships among ZEB2, MEIS1, MYB, and MEF2C (Fig. 1G, 1H), which were also confirmed at the protein level (Fig. 1I). Due to the absence of CP-CML cell lines, we collected bone marrow mononuclear cells from chronic phase patients and found that such regulatory interactions as observed in BP-CML cells did not exist in chronic phase CMLs (Fig. 1J). Consistent with our prediction, our co-immunoprecipitation assays further confirmed the interaction dynamics among these four TFs—specifically, any single TF can stably interact with the other three TFs (Fig.1K).

### 2. The TFs of the circuit mutually activate the transcription of each other

After identifying that MEF2C, MEIS1, MYB, and ZEB2 form a transcriptional regulatory circuit, we next explored the regulatory mechanisms among these four TFs. In two BP-CML cell lines, K562 and LAMA84, we performed CUT&Tag assays using their respective antibodies and antibodies targeting H3K4me1 (active/poised enhancers), H3K4me3 (active promoters), and H3K27ac (active enhancers or promoters). The results revealed that all four TFs co-binding to the promoters or enhancers of their own loci and the other three TFs in both cell lines (Fig. 2A-2B, S2A-S2C), suggesting they may directly transcriptionally activate their own expression and that of each other. To validate their co-binding at the same enhancer or promoter regions, we scanned their motifs at the co-bound loci and found that the distances between the motifs of the four TFs were mostly within one nucleosome (247 kb)^24^ (Fig. 2C, S2D-S2F).

**Figure 2.**
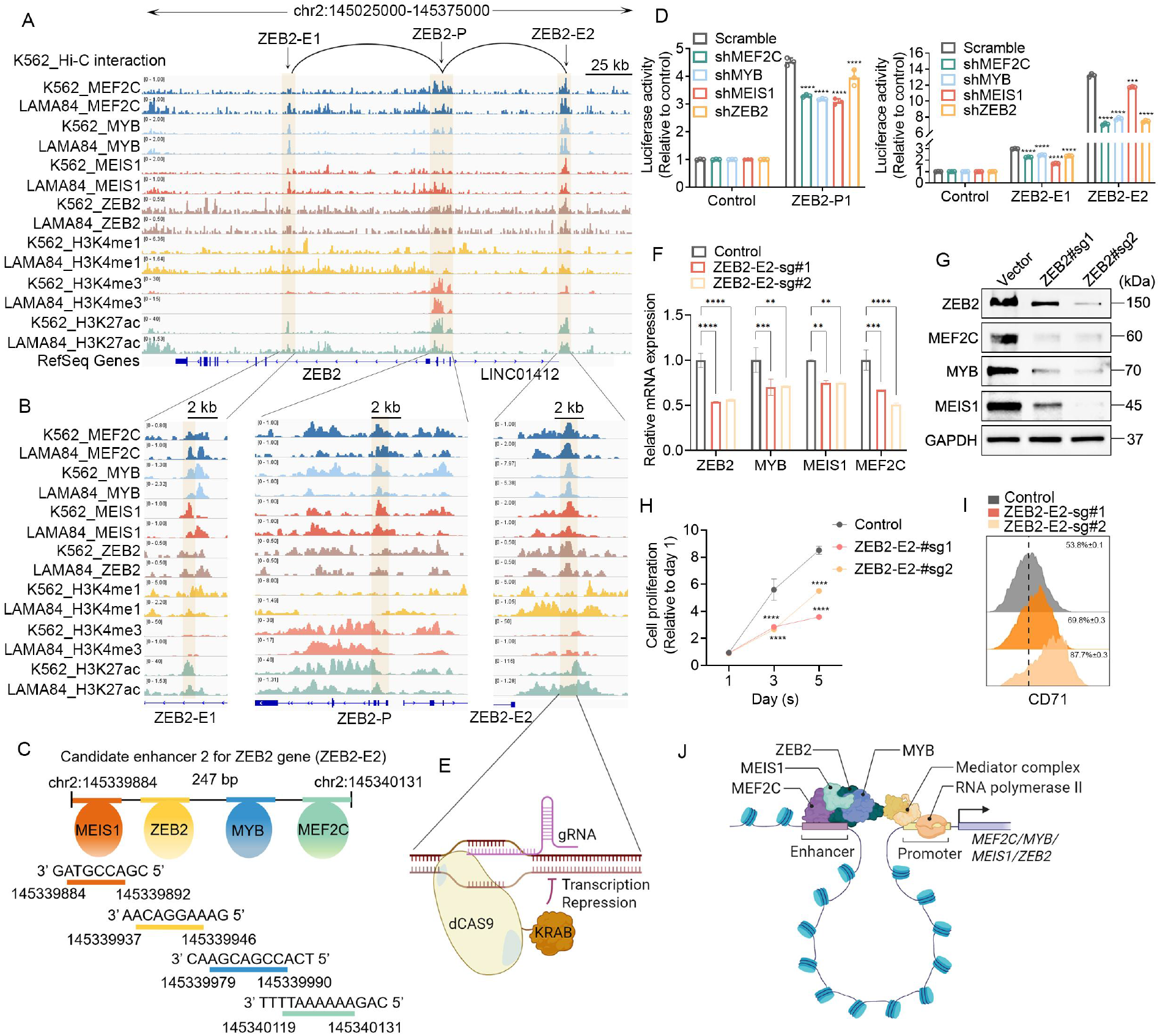
MEF2C, MYB, MEIS1, and ZEB2 co-bind to the promoter or enhancer regions of ZEB2 to activate its transcription. A,B. The tracks show profiles of ChIP-Seq or CUT&Tag signals of indicated antibodies at ZEB2 gene loci. C. A schematic representation of the nearest distance pattern of MEF2C, MYB, MEIS1, and ZEB2 motifs at ZEB2-E2. D. Promoter and enhancer activities were measured by luciferase reporter assays in K562 cells in either the presence or absence of knockdown indicated TF. E, This figure illustrates the working mechanism of Krab-dCas9. F,G. The mRNA and protein level were detected in K562 cells with control or ZEB2-E2-sg transfection. H, I. Cell proliferation and defferentiation were measured under control or ZEB2-E2-sg transfection. J. Schematic diagram of the reciprocal regulatory network composed by MEF2C, MYB, MEIS1, and ZEB2.

Taking the ZEB2 gene locus as an verify, we validated the transcriptional regulatory effects of this loop. The promoter (ZEB2-P) co-binding by the four TFs was cloned into the pGL3-Basic dual-luciferase reporter vector, while the two enhancer regions (ZEB2-E1 and ZEB2-E2) were separately cloned into the pGL3-promoter dual-luciferase reporter vector. The results showed that ZEB2-P, ZEB2-E1, and ZEB2-E2 all exhibited significant transcriptional activation activity. Notably, knockdown of each TFs suppressed the transcriptional activity of ZEB2-P, ZEB2-E1, and ZEB2-E2 (Fig. 2D). These findings indicate that the four TFs regulate ZEB2 transcription by modulating the transcriptional activity of the ZEB2-P, ZEB2-E1, and ZEB2-E2 regions. Furthermore, we designed sgRNAs targeting the ZEB2-E2 locus and cloned them into the KRAB-deadCas9 vector. After inhibiting the transcriptional activity of ZEB2-E2, the mRNA and protein levels of the four TFs were all downregulated (Fig. 2E-2G). Concurrently, cell proliferation was inhibited (Fig. 2H), and the differentiation capacity of the cells was enhanced (Fig. 2I). This finding strongly supports the conclusion that MEF2C, MEIS1, MYB, and ZEB2 form a multi-subunit complex to achieve spatial co-localization within the same enhancer or promoter region. Through this coordinated mechanism, they synergistically activate their own transcription and that of each other (Fig. 2J). This complex-mediated transcriptional regulation model provides critical experimental evidence for deciphering the molecular mechanisms underlying epigenetic reprogramming during the blast crisis phase of CML.

### 3. The TFs cooperatively regulate global chromatin accessibility in blast CML cells

We have confirmed that MEF2C, MEIS1, MYB, and ZEB2 form a BP-CML-specific transcriptional regulatory circuit, and have further elucidated the regulatory mechanisms underlying their mutual interactions. To explore whether the circuit is capable of modulating chromatin accessibility to regulate global gene expression network, we conducted a classification analysis of CUT&Tag data from the two cell lines based on TFs binding profiles. The results revealed that 74.6% of promoters and 28.2% of enhancers were bound by two or more TFs in K562 cells, and this co-bind phenomenon also consistant in LAMA84 cells (Fig. 3A-3B, S3A). Notably, across both promoter and enhancer regions, co-binding sites of the four TFs (quard group) exhibited the strongest H3K27ac signals (Fig. 3C, S3B). These co-binding regions accounted for a relatively higher proportion of SEs (Fig. 3D, S3C), and correspondingly, the expression of these co-binding sites nearby genes was also significantly higher than other groups (Fig. 3E, S3D). These findings suggest that MEF2C, MYB, MEIS1, and ZEB2 synergistically promote the transcriptional activation of downstream target genes.

**Figure 3.**
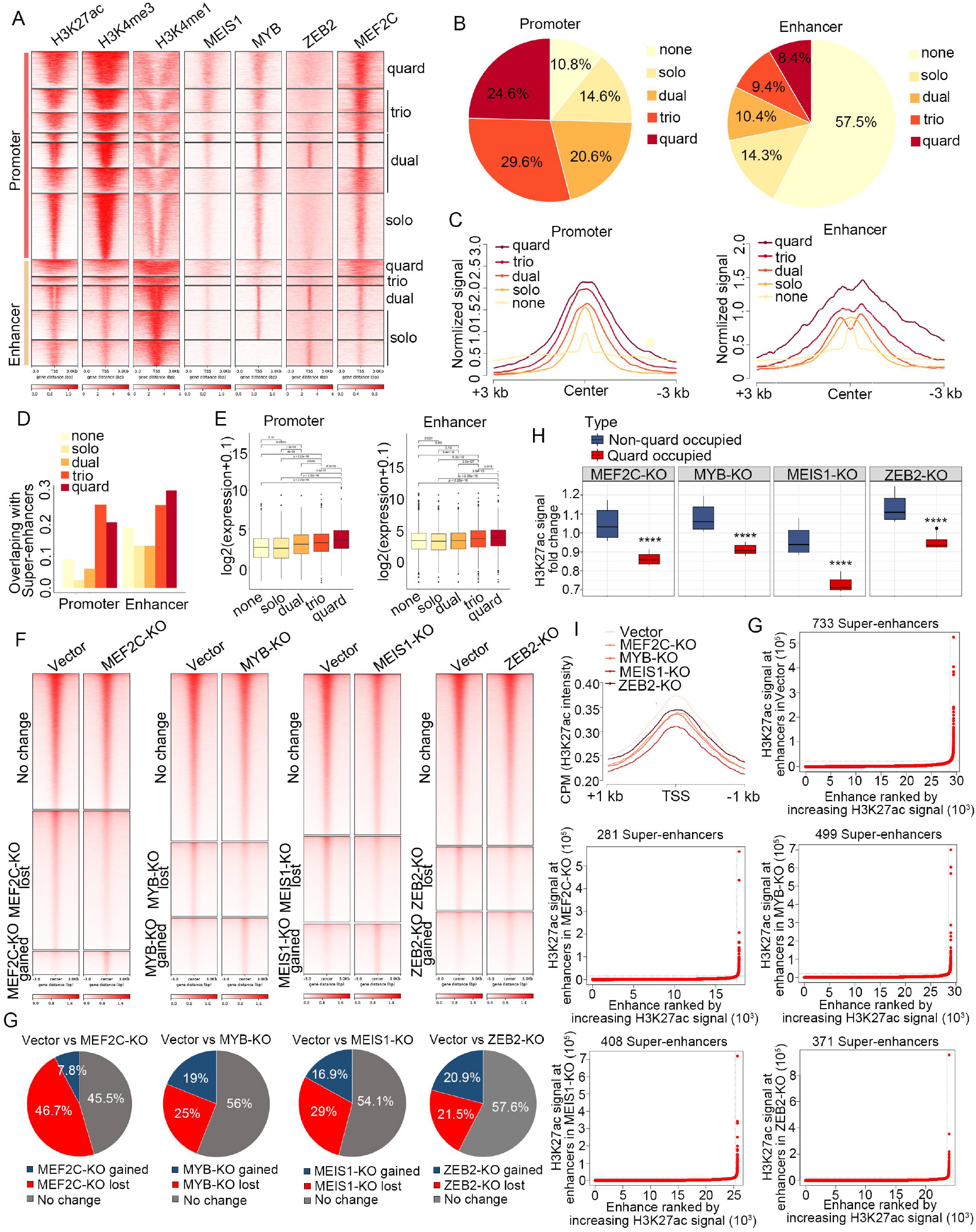
This circuit regulates the chromatin accessibility landscape in BP-CML cells. A. Heatmaps of CUT&Tag or ChIP-Seq signals of indicated factors in K562 cells, stratified by different combinatorial binding patterns. B. The pie chart show the co-binding overview of MEF2C, MYB, MEIS1, and ZEB2 at promoters and enhancers. C. The normalized signal intensity of H3K27ac among the peaks of indicated groups. D. The proportion of super enhancers (SEs) among peaks in the indicated groups. E. The mRNA expression of adjacent to the indicated peaks in K562 cells. F. H3K27ac-ChIP-Seq heatmaps of differential accessible regions between scramble and each TF-knockout in K562 cells. G. The pie chart show the proportion of gained or lost peaks under TF knockout. H. Changes in H3K27ac signals at regions with or without co-binding of the four TFs (TF-KO vs Vector). I. H3K27ac intensity of BP-CML gained peaks in indicated groups. J. ROSE map of identified SE in indicated groups.

We hypothesized that the TFs co-binding at shared chromatin regions likely regulates target gene transcription via alterations in chromatin accessibility. To test this, we conducted H3K27ac CUT&Tag assays after individually knocking out the MEF2C, MYB, MEIS1, and ZEB2 to measure changes in chromatin accessibility. Consistent with the results of TF knockdown, the knockout of any single TF significantly decreased the expression levels of the other three TFs (Fig. S3E). Compared to Vector, TF knockout reduced chromatin accessibility in >25% of regions; notably, MEF2C knockout led to a 46.7% decrease in H3K27ac signals at these regions (Fig. 3F and 3G). Importantly, depleting any component of the regulatory circuit significantly diminished chromatin accessibility signals at regions co-bound by these four TFs (Fig. 3H). Intriguingly, TF knockout also reduced H3K27ac signals at BP-CML gained peak regions (Fig. 3I) and markedly decreased the number of super enhancers (SEs) in BP-CML cells (Fig. 3G).

### 4. The factors cooperatively orchestrate the transcriptional network of blast CML cells

Our results confirmed that MEF2C, MEIS1, MYB, and ZEB2 coordinately regulate chromatin accessibility in BP-CML cells. To determine whether this ultimately led to transcriptomic remodeling in BP-CML cells, we performed RNA sequencing following individual knockdown of each TF in K562 cells. GSEA enrichment analysis revealed that the down-regulated gene sets induced by knockdown of each TF exhibited significant similarity, so as the up-regulated gene sets (Fig. 4A-4B, S3F), indicating that the four TFs display remarkably consistent regulatory patterns in controlling transcriptional programs. Importantly, the co-downregulated genes following knockdown of each TF was significantly enriched in the gene sets upregulated in BP-CML across both RNA-Seq datasets (Fig. 4C, S3G). Chromatin accessibility of the 302 co-downregulated genes was significantly suppressed following TF knockout (Fig. 4D). Among these, RUNX1—representative of the 302 genes—exhibited co-binding of MEF2C, MEIS1, MYB, and ZEB2 at its enhancer. Upon TF knockout, H3K27ac signals at the RUNX1 enhancer loci were reduced, concomitant with decreased RUNX1 mRNA expression (Fig. 4E). These results collectively indicate that MEF2C, MEIS1, MYB, and ZEB2 co-binding at enhancer regions of downstream genes enhances chromatin accessibility at these enhancers, thereby promoting gene expression.

**Figure 4.**
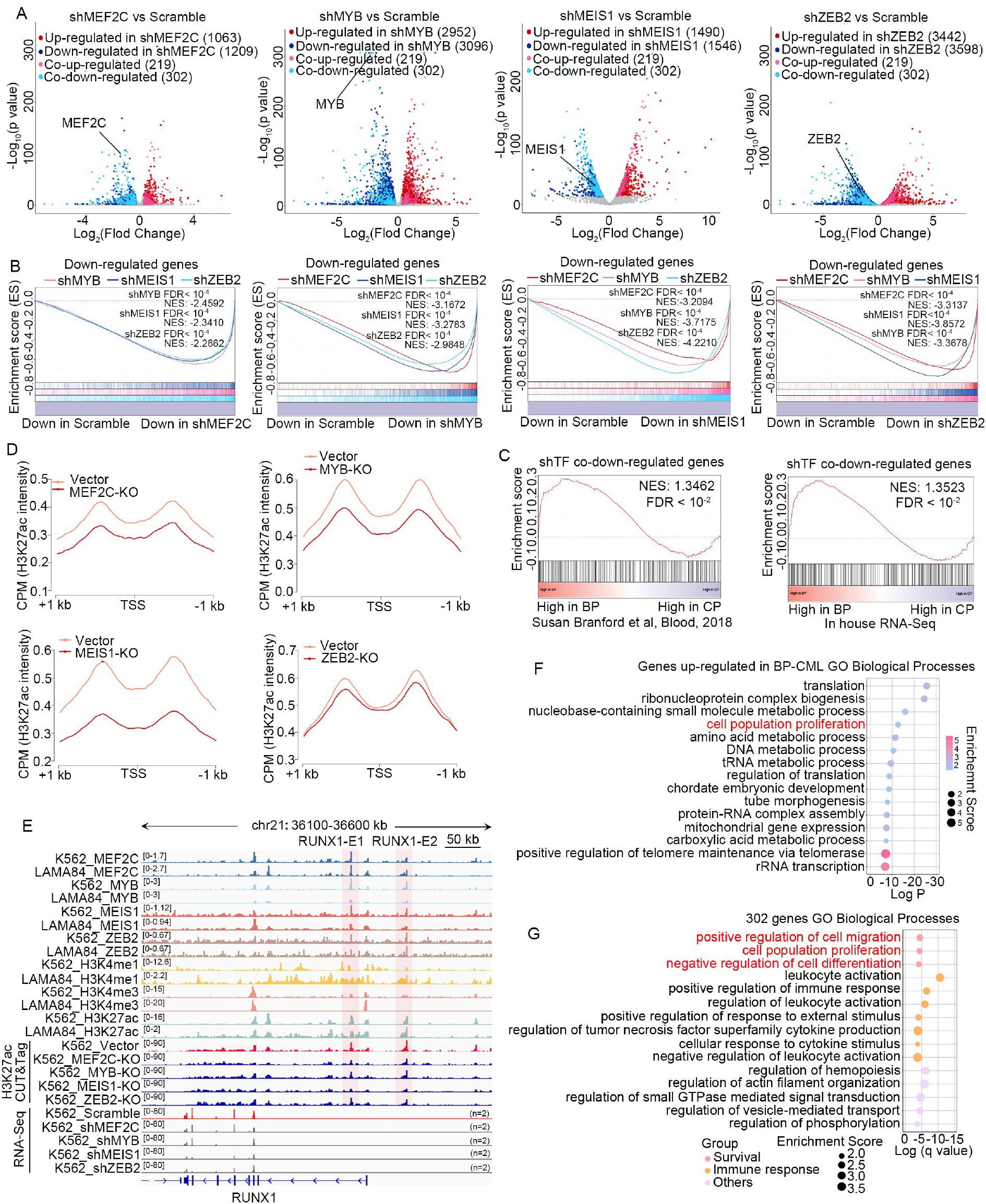
MEF2C, MYB, MEIS1, and ZEB2 cooperatively regulate the transcription of downstream genes. A. Volcano plot of mRNA expression changes following transcription factor (TF) knockdown. Up regulated in shTFs (MEF2C, MYB, MEIS1, ZEB2), log2 fold-change>0.584, p-value<0.05; down regulated in shTFs (MEF2C, MYB, MEIS1, ZEB2), log2 fold-change<-0.584 and p-value<0.05. B. Genes down-regulated by knocking down each TF are subjected to GSEA enrichment analysis with the RNA-Seq data of the other three knockdown. C. Genes co-down-regulated by knocking down MEF2C, MYB, MEIS1, ZEB2 are subjected to GSEA enrichment analysis with the RNA-Seq data of BP and CP-CMLs. D. The H3K27ac intensity of co-down-regulated genes’ promoter between Scramble and TF knocking out. E. The IGV tracks of indicated factors shown at RUNX1 loci. F. GO biological enrichment of genes up-regulated in BP-CML. G. GO biological enrichment of genes co-down-regulated in MEF2C, MYB, MEIS1and ZEB2 knockdown.

To investigate how MEF2C, MEIS1, MYB, and ZEB2 modulates the malignant phenotypes of BP-CML cells, we first performed Gene Ontology (GO) pathway enrichment analysis on the intersection of BP-CML up-regulated genes identified in both RNA-Seq datasets. The results revealed significant enrichment in cell proliferation pathway (Fig. 4F), indicating enhanced proliferative capacity during disease progression from chronic phase to blast crisis. We then conducted GO pathway enrichment analysis on gene sets that were commonly down-regulated or up-regulated following knockdown of the four TFs. The results illustrated that cell differentiation and cell proliferation pathways were significantly enriched (Fig. 4G, S3H), which further suggested that MEF2C, MEIS1, MYB, and ZEB2 may promote proliferation and suppress differentiation of BP-CML cells. This finding is consistent with the clinical observations that during disease progression from chronic phase to blast crisis, tumor cells display augmented proliferation and compromised differentiation.

### 5. Silencing the four TFs inhibits the survival of blast CML cells

To validate the functional roles of MEF2C, MEIS1, MYB, and ZEB2 in BP-CML cells, we individually knocked down TFs in BP-CML cells, this resulted in significantly inhibited cell proliferation and clonogenic capacity (Fig. 5A, S4A). Notably, the knockdown of TFs also led to enhanced myeloid differentiation capacity, including megakaryocytic, erythroid, and granulocytic lineages (Fig. 5B-5D, S4B-S4C). Given that these TFs synergistically drive the malignant phenotypes of blast phase (BP) cells, we conducted a preliminary exploration of their cooperative interactions. Our results demonstrated that, compared with knockdown of a single TF, simultaneous knockdown of two TFs significantly enhanced both the inhibition of cell proliferation and the activation of cellular differentiation (Fig. S4D-S4E).

**Figure 5.**
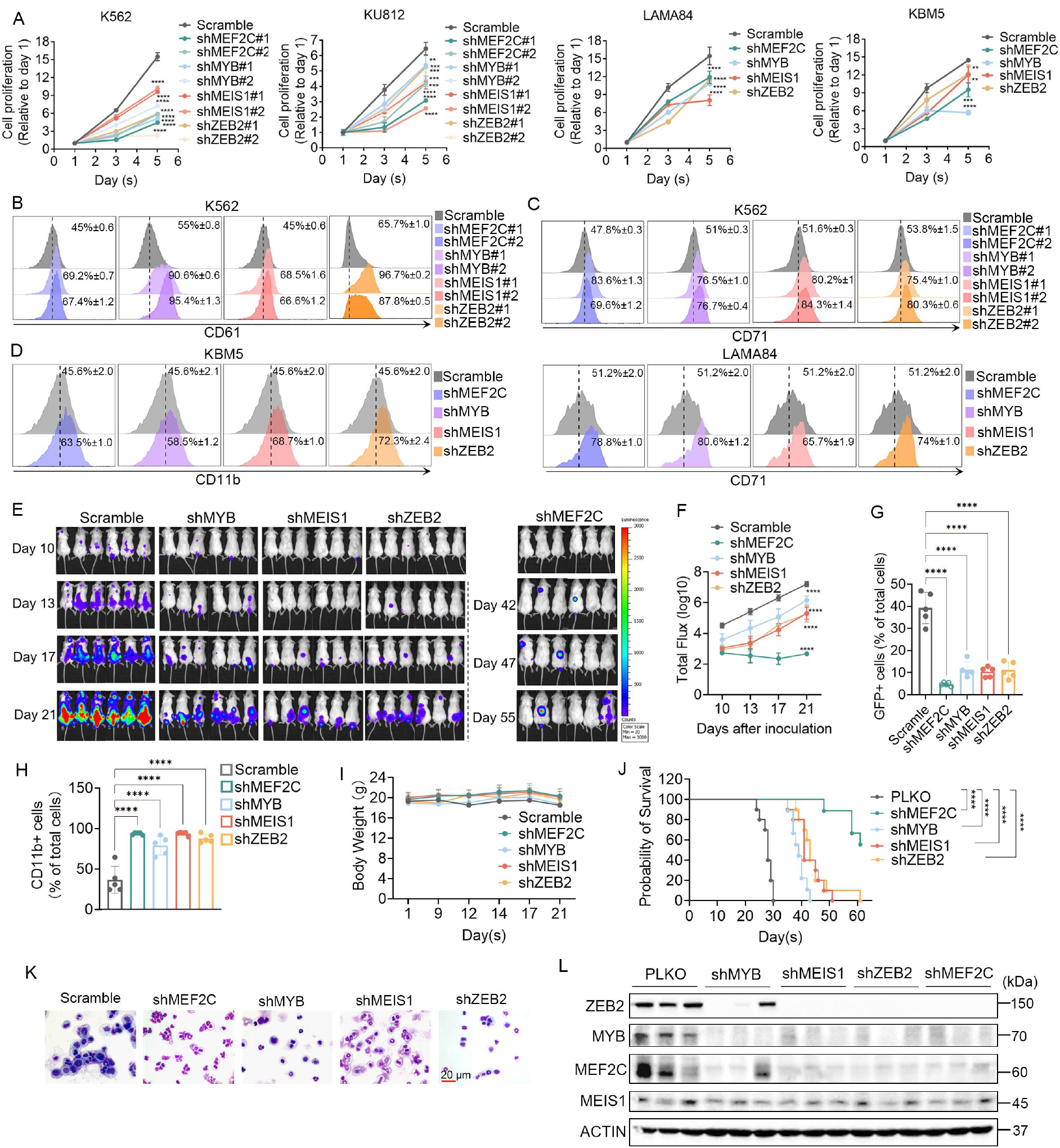
MEF2C, MYB, MEIS1, and ZEB2 knockdown impair the malignancy of BP-CML both *in vitro and in vivo*. A. Cell viability were measured under TF knockdown on indicated days in BP-CML cell lines. Cell differentiation were detected by flow cytometer with TF knockdown, (B) CD61 for megakaryocyte differentiation (1 nM PMA treatment 24h), (C) CD71 for erythroid differentiation (10 μM hemin treatment 48h), and (D) CD11b for granulocyte differentiation (1 μM ARTR treatment 72h). E. Bioluminescent image of NSG mice at indicated days after implantation with the Scramble or TF-knockdown KBM5 cells, respectively (n=6). F. The statistical chart corresponding to Figure. E. The proportion of GFP positive cells (G) and CD11b positive cells (H) in bone marrow cells isolated from NSG mice (n=5). I. The body weight of indicated group NSG mice. J. Kaplan-Meier survival curves of NSG mice in indicated groups (n = 10). K. The represent image shown that Gimsa staining were performed in bone marrow cells isolated from indicated NSG mice (n=3). L. Weston Blot were performed to measure the protein level of bone marrow cells isolated from NSG mice (n=3).

Consistent with prior studies, tail vein injection of KBM5 cells into immunocompromised mice reliably establishes a BP-CML mouse model^25, 26^. To validate the in vivo efficacy of the TFs knockdown in suppressing BP-CML progression, we generated luciferase-expressing KBM5 cells within stably express either a GFP-tagged empty vector or shRNA plasmids targeting CRC-TFs, followed by tail vein inoculation. Our results demonstrated that silence the TFs significantly suppressed BP-CML progression (Fig. 5E, 5F), as evidenced by reduced GFP-positive tumor cell proportion (Fig. 5G), increased granulocytic CD11b-positive cell percentage (Fig. 5H), and no significant changes in mouse body weight (Fig. 5I). Notably, survival time of BP-CML mice was significantly prolonged (Fig. 5J). Giemsa staining of bone marrow cells revealed a marked reduction in primitive cells in the TF-knockdown group compared with controls (Fig. 5K). Western blot analysis of isolated bone marrow cells confirmed that the TFs maintained their mutual regulatory interactions (Fig. 5L). Collectively, these findings demonstrate that MEF2C, MEIS1, MYB, and ZEB2 sustain the mutual regulatory pattern in vivo. Knockdown of these TFs effectively inhibits BP-CML cell proliferation, promotes cellular differentiation both *in vitro* and *in vivo*.

### 6. The circuit activates purine metabolism and Rho GTPase signaling pathway

We investigated TF-driven mechanisms of CML blast crisis. Pathway enrichment analysis of upregulated genes in two CML cohorts (in-house and Branford et al. RNA-Seq) revealed strong overlap with cell proliferation, hematopoietic lineage, purine metabolism, amino acid metabolism, and RHO GTPase signaling pathways (Fig. S5A). Intersection of TF-bound genes (K562 cells) and BP-CML upregulated genes identified nucleotide metabolism and RHO GTPase pathways as critical for blast progression (Fig. 6A and 6B). Co-binding genes, ranked by TF knockdown effects (Fig. 6C and 6D), revealed downstream mediators of TF regulation. We identification of the nucleotide metabolism-related top-ranked GLUL. GLUL (glutamine synthetase) is a key metabolic enzyme that mediates the synthesis of glutamine from glutamate within cells. Glutamine serves as a critical nitrogen source for the biosynthesis of purine and pyrimidine nucleotides. Interestingly, GTP serves as both an intermediate product in purine nucleotide synthesis and a critical molecule for activating RHO GTPase signaling. Studies have demonstrated that alterations in purine nucleotide metabolism and glutamine synthetase mediate the activation of RHO GTPase signaling ^27, 28^. Based on this, we hypothesize that the TFs may promote blast crisis progression by regulating GLUL to simultaneously enhance purine nucleotide synthesis and activate RHO GTPase signaling.

**Figure 6.**
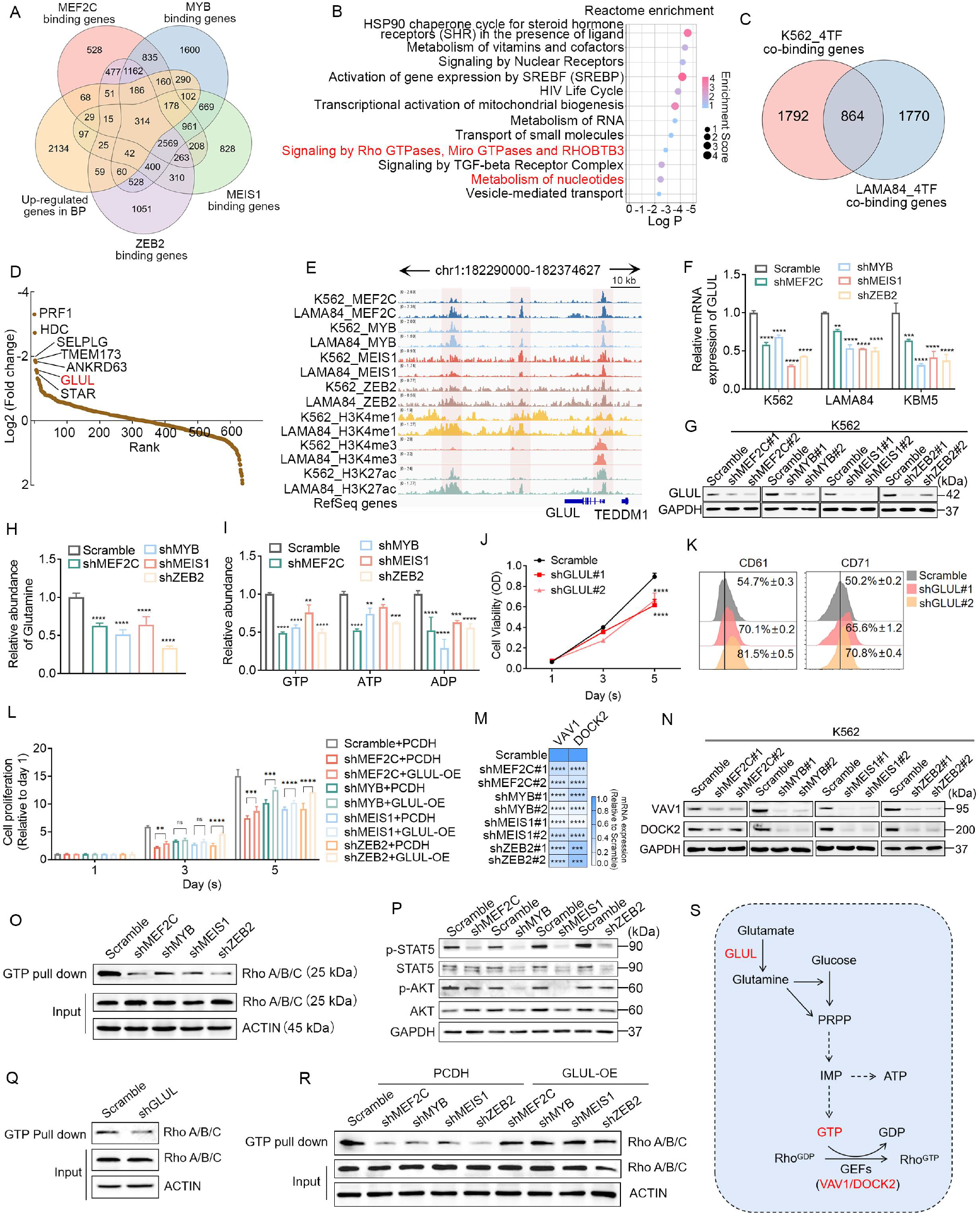
The circuit activates purine metabolism and Rho GTPase signaling pathway. A. The overlap genes of binding by MEF2C, MYB, MEIS1, ZEB2 and up-regulated in BP-CML. B. The pathway enrichment of 314 overlapping genes (Figure A) (P< 0.01). C. The overlap genes of MEF2C, MYB, MEIS1, and ZEB2 co-binding genes in K562 and LAMA84 cells. D. Ranking of 864 genes by log2(Fold change) after MEF2C, MYB, MEIS1, and ZEB2 knockdown. E. The tracks show profiles of ChIP-Seq or CUT&Tag signals of indicated antibodies at GLUL gene loci. Relative mRNA (F) and protein (G) expression of GLUL under MEF2C, MYB, MEIS1, and ZEB2 knockdown. H. The abundance of Glutamine measured by ELISA assay with TF knockdown and Scramble. I. The abundance of GTP, ATP, and ADP measured by HLPC with TF knockdown and Scramble. J. Cell viability were measured under GLUL knockdown on indicated days in K562 cells. K. Cell differentiation were measured under GLUL knockdown in K562 cells. L. Cell viability was evaluated in K562 cells with TF knockdown and Scramble, followed by lentiviral overexpression of PCDH or GLUL. The mRNA (M) and protein (N) expression of VAV1 and DOCK2 under MEF2C, MYB, MEIS1, and ZEB2 knockdown. O. GTP pull down performed with MEF2C, MYB, MEIS1, and ZEB2 knockdown in K562 cells. P. The protein level of indicated factors were detected under TFs silencing. Q. GTP pull down performed with GLUL knockdown in K562 cells. R. GTP pull down performed in K562 cells with TF knockdown and Scramble, followed by lentiviral overexpression of PCDH or GLUL. S. Schematic diagram of crosstalk between purine metabolism and Rho GTPase signaling pathway.

CUT&Tag assays revealed co-localization of MEF2C, MEIS1, MYB, and ZEB2 at both enhancers and promoters of GLUL across two BP-CML cell lines (Fig. 6E). Moreover, we detected a higher mRNA expression of GLUL in BP-CML compared to CP-CML (Fig.S5B). These results strongly suggest their synergistic contribution to GLUL transcriptional activation in BP-CML. Consistent with this, knockdown of the TFs in BP-CML cells led to concurrent downregulation of GLUL mRNA and protein levels (Fig. 6F, 6G), diminished intracellular glutamine content (Fig. 6H), and reduced levels of key purine nucleotide intermediates (including GTP, ADP, and ATP) (Fig. 6I). Disruption of nucleotide synthesis further triggered G1-phase cell cycle arrest (Fig. S5C), culminating in inhibited cell proliferation. Notably, knockdown of GLUL in BP-CML cells recapitulated these phenotypic effects: it reduced intracellular glutamine and purine nucleotide intermediates (GTP, ADP, ATP), suppressed proliferation, and enhanced differentiation capacity—observations mirroring those induced by TF knockdown (Fig. S5D-S5F, 6J-6K). Conversely, GLUL overexpression partially rescued the decreases in glutamine and purine metabolites, as well as the proliferation inhibition caused by TF knockdown (Fig. S5G-S5H, Fig. 6L). Collectively, these results demonstrate that MEF2C, MEIS1, MYB, and ZEB2 drive purine nucleotide synthesis through their activation of GLUL transcription.

Rho GTPase is a family of critical cell signaling proteins that exert essential regulatory roles in cell proliferation, differentiation, and motility^29^. Rho GTPase primarily comprising three subfamilies: RhoA/B/C, Rac1/2/3, and CDC42. Their activation—defined as the exchange of GDP for GTP—is catalyzed by GEFs (guanine nucleotide exchange factors). Our study revealed that MEF2C, MEIS1, MYB, and ZEB2 may directly drive the expression of GEF enzymes VAV1 and DOCK2 (Fig. S5I). Subsequent validation in BP-CML cells confirmed that TF knockdown suppressed both mRNA and protein expression of VAV1 and DOCK2 (Fig. 6M, 6N). We examined the activation of RhoA/B/C (highly expressed in BP-CML cells) using GTP pull-down assays. Knockdown of the TFs reduced GTP-bound RhoA/B/C levels (Fig. 6O), concomitant with inhibited phosphorylation of downstream AKT and STAT5 (Fig. 6P), indicating suppression of Rho GTPase signaling. These results collectively demonstrate that the TFs mediate Rho GTPase activation through transcriptional upregulation of GEFs VAV1 and DOCK2. Building on our prior hypothesis that TFs promote GTP synthesis and subsequent Rho GTPase activation by up-regulate GLUL, we validated this axis by knocking down GLUL in BP-CML cells. Consistent with TF knockdown phenotypes, GLUL depletion reduced GTP-bound RhoA/B/C levels and suppressed AKT/STAT5 phosphorylation (Fig. 6Q, S5J). Notably, GLUL overexpression partially rescued the inhibitory effects of TF knockdown on RhoA/B/C activation (Fig. 6R). Given that sustained activation of BCR-ABL kinase activity is critical for maintaining the survival of CML cells, we investigated whether MEF2C, MYB, MEIS1, and ZEB2 regulate BCR-ABL activity. Our results demonstrated that TFs do not modulate BCR-ABL expression or its kinase activity (Fig. S5J), indicating that the TF-mediated regulatory circuit exerts its effects on CML cell survival independently of the BCR-ABL pathway.

Collectively, these findings support three key conclusions (Fig. 6S): 1) TFs transcriptionally activate GLUL to enhance purine synthesis, thereby promoting cell proliferation; 2) TF-driven activation of the purine metabolism pathway increases GTP production, leading to Rho GTPase signaling activation; 3) TFs transcriptionally upregulate GEF enzymes VAV1 and DOCK2 to potentiate Rho GTPase signaling.

### 7. Mebendazole restrains the malignancy of blast CML

To explore the clinical potention of targeting the transcriptional regulation loop, we try to found inhibitors against the factor of this circuit. Emerging evidence demonstrates that the clinically anthelmintic drug Mebendazole (MBZ) which can degrade MYB protein^30^, and the non-directly targeted MEF2C inhibitor YKL-05-099 both exert therapeutic effects on hematological malignancies, including AML^31^. The mRNA and protein level of MEF2C, MYB, MEIS1, and ZEB2 were concomitantly diminished in BP-CML cells with MBZ or YKL-05-099 treatement (Fig. 7A-7B and S6A-S6C). MBZ and YKL-05-099 both exhibited the inhibition of cell viability in BP-CML cell lines (Fig. 7C and S7D). Considering the clinical translational potential, we prioritize MBZ as the candidate drug for further investigation, as it has already applied on clinical and demonstrated significant inhibitory effects against leukemia in vitro. As expected, MBZ exhibited anti-leukemia activity against bone marrow mononuclear cells from CML patients without toxicity on the PBMCs from healthy donors (Fig. 7D). MBZ treatment also induced the cell differentiation of BP-CML cells (Fig. 7E).

**Figure 7.**
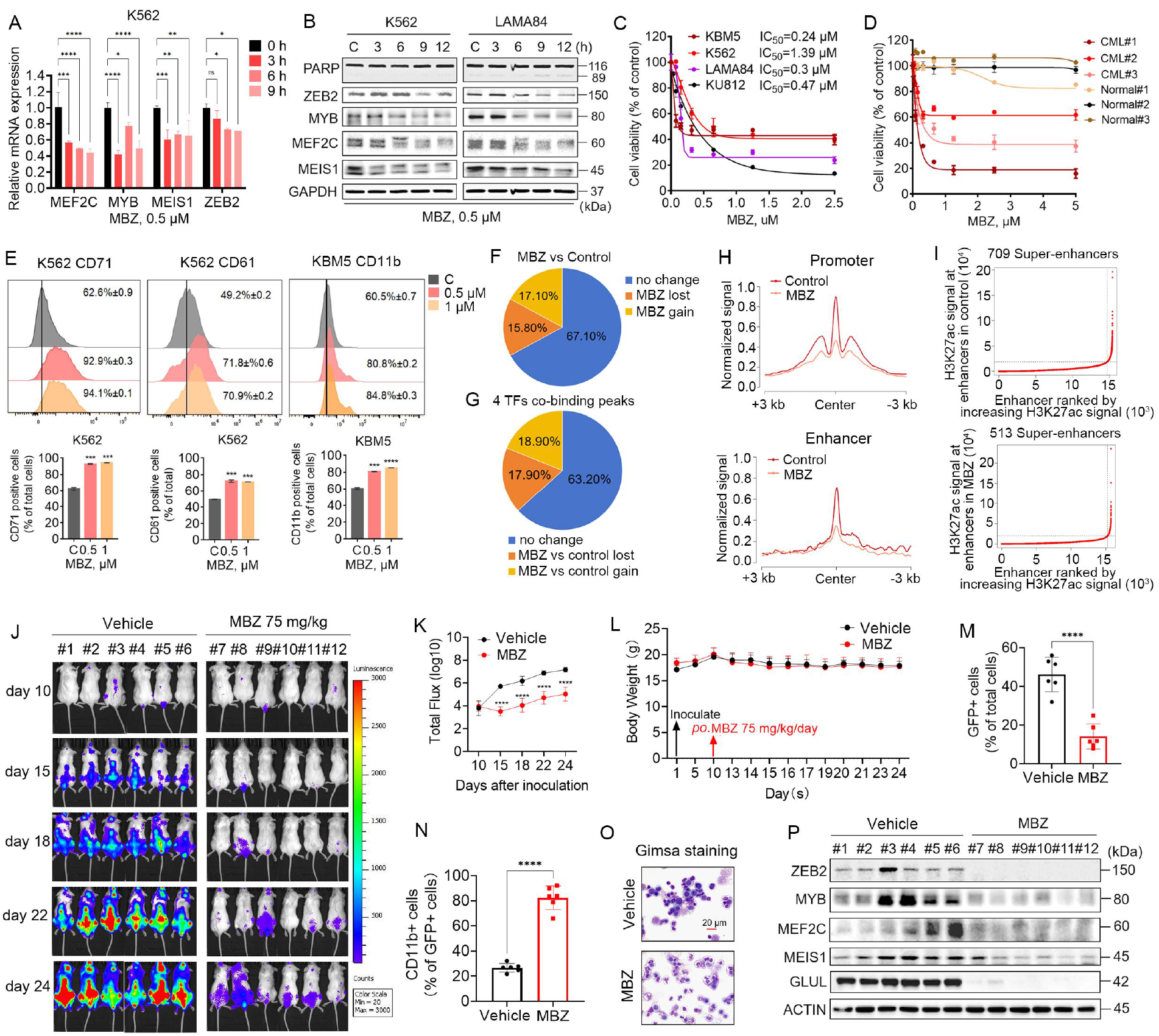
Mebendazole impair the malignancy of BP-CML both *in vitro and in vivo*. A. K562 cells were treated with Mebendazole, and then the indicated mRNA were detected byRT-qPCR. B. K562 and LAMA84 cells were treated with Mebendazole, and then the indicated proteins were detected by Western blots. C. Treatment of CML cells (KBM5, K562, LAMA-84, KU812) with Mebendazole, MTS assay performed to measure cell viability after 72 h. D. Treatment of PBMC from CML patients or healthy donor with Mebendazole, MTS assay performed to measure cell viability after 72 h. E. Representative flow cytometry histogram of surface CD71, CD61 or CD11b expression in K562 or KBM5 cells treated with Mebendazole. F. Pie chart illustrating altered proportion of H3K27ac peaks upon given of Mebendazole in K562 cells. G. Pie chart illustrating altered proportion of four TFs co-binding peaks upon given of Mebendazole in K562 cells. H. Line plots of H3K27ac ChIP-Seq signals from indicated groups of peaks in K562 cells. I. Hockey-stick plots displaying the altered number of SE-related genes in K562 cells treated with Mebendazole. J. Bioluminescence imaging of luciferase-KBM5-xenotransplanted mice after treatment with Mebendazole (75mg/kg). K. Statistics of bioluminescence imaging in mice after Mebendazole treatment. Data were derived from panel (J). L. Mouse weight measurements were performed during daily treatment with 75mg/kg Mebendazole (n=10). M. Percentages of GFP positive cells in bone marrow. N. Percentages of CD11b positive leukemia cells in bone marrow. O. The represent image shown that Gimsa staining were performed in bone marrow cells isolated from indicated NSG mice (n=3). P. Indicated protein expression in bone marrow cells of control (n=6) and Mebendazole treatment (n = 6) were detected by Western blots.

We next sought to understand the role of MBZ in chromatin accessibility by performing H3K27ac ChIP-seq in either the presence or absence of MBZ. MBZ altered the H3K27ac signal across thousands of genomic loci (15.8% of lost peaks and 17.1% of gained peaks) (Fig. 7F and S7E). MBZ had a more profound impact on its direct target MYB-binding peaks, although there is also a certain overlap in the binding peaks for the other three TFs respectively (Fig. S7F). In contrast, MBZ treatment selectively modulated chromatin accessibility at regions co-occupied by four TFs (17.9% of lost and 18.9% of gained) (Fig. 7G). MBZ treatment reduced the chromatin accessibility in the promoter and enhancer regions (Fig. 7H). Moreover, MBZ treatment reduced the number of SEs compared with control (Fig. 7I). These results suggest that MBZ extensively suppress the chromatin accessibility which consistent with TFs knockdown in BP-CML cells.

Oral administration of MBZ initiated at day 10 post-KBM5-luciferase engraftment in NSG mice, was found to have significant anti-leukemic efficacy with stable body weight trajectory and toxicity indicators in serum (Fig. 7J-7L, S7G), demonstrating its potential as a safe and effective therapeutic agent for CML blast crisis. Futhermore, MBZ administration significantly reduced peripheral leukocytosis and attenuated leukemic bone marrow infiltration while inducing CD11b^+^ granulocytic maturation in residual leukemic populations (Fig. 7M-7O, S7H). The protein level of MEF2C, MYB, MEIS1, ZEB2 and downstream transcriptional targets were decreased with MBZ treatment (Fig.7P).

## Disscussion

Cell-type-specific transcriptional regulatory programs are typically orchestrated by transcriptional networks driven by super-enhancers^32^. Disrupting super-enhancer-mediated transcriptional programs has emerged as a pivotal therapeutic strategy in cancer^33^. In this study, we collected CML specimens from chronic and blast phases, performed transcriptomic and epigenetic modification high-throughput sequencing, and validated findings using cell lines and patient-derived bone marrow cells. This approach identified a CML blast-specific transcriptional regulatory circuit comprising MEF2C, MYB, MEIS1, and ZEB2. Notably, these transcription factors exhibit three key characteristics: 1) co-binding to each other’s promoters or enhancers to reciprocally regulate transcription; 2) direct protein-protein interactions; 3) collaborative reprogramming of chromatin accessibility and transcriptomic landscapes in blast cells. Our work first demonstrates that this MEF2C-MYB-MEIS1-ZEB2 circuit serves as a central driver of CML blast crisis. Functional validation revealed that silencing components of this circuit significantly suppressed CML blast progression *in vitro* and *in vivo*. Although MYB and MEIS1 have been reported to promote CML blast transformation, our study innovatively identifies their synergistic roles within this circuit rather than acting independently, as revealed through patient-derived tumor sample screening and high-throughput approaches. Recent studies suggest that high concentrations of TFs within active enhancer regions depend on liquid-liquid phase separation condensates^34^. We hypothesize that the identified circuit components likely exhibit phase separation properties as well, given their incorporation of multiple intrinsically disordered region (IDR)-containing sequences. This hypothesis merits further investigation as a promising direction for elucidating the mechanistic basis of their transcriptional regulatory functions.

Mechanistically, we uncovered that this circuit activates purine metabolism and Rho GTPase signaling pathways through transcriptional activation of GLUL (catalyzing the biosynthesis of glutamate to glutamine) and GEF enzymes. Prior studies demonstrated that branched-chain amino acids (BCAAs) are metabolized by Bcat1 into branched-chain α-keto acids (BCKAs), with BCAA-derived amino groups transferred to α-ketoglutarate (α-KG) to generate glutamate^35^. While BCAA levels are elevated in blast-phase CML compared to chronic phase, glutamate levels remain unchanged. Our findings reveal that hyperactive nucleotide metabolism in blast cells, driven by circuit overexpression, requires GLUL to convert excess glutamate into glutamine as a metabolic precursor—a plausible explanation for stabilized glutamate levels.

Given the circuit’s critical role in CML blast progression, targeting this network represents a promising therapeutic avenue. Screening clinically approved agents, we focused on mebendazole (MBZ), previously shown to destabilize MYB protein in acute myeloid leukemia (AML). Our results confirm that MBZ suppresses circuit component expression and effectively mitigates CML blast progression in vitroand in vivo. While clinical efficacy in CML remains unvalidated, MBZ’s established safety profile and mechanistic activity position it as a high-potential translational candidate, warranting further clinical investigation.

## Acknowledgments

This work was supported by the National Natural Science Foundation of China (82170177/H0809), the Natural Science Foundation of Guangdong Province (2023A1515011976) to XPS; National Natural Science Foundation of China (82303536), the Natural Science Foundation of Guangdong Province (2025A1515010451) to LLJ; Guangdong Basic and Applied Basic Research Foundation (2024A1515010703), National Natural Science Foundation of China (32200538) to YYZ.

## Authorship

Contribution: XPS, YYZ and LLJ conceived and directed the study. LLJ, EZL and JXF performed the experiments. LZJ, GJP, BYL, and QM provided bioinformatics analysis. MY, HCZ, and ACL provided statistical and technical support. JBL, PLL, JHC, and LB were responsible for providing clinical advice. XPS, LLJ, and YYZ provided funding and supervision. LLJ, EZL, and JXF wrote the first draft of the manuscript. XPS and YYZ revised the manuscript. All the authors had full access to the data, and reviewed the manuscript before submission.

## Data availability

The sequencing data generated in this study were submitted to Arrayexpress Database under accession numbers E-MTAB-16036 (ChIP-Seq targeting H3K27ac in Bone Marrow cells from CML patients), E-MTAB-16052 (ChIP-Seq and CUT&Tag targeting histone markers or TFs in BP-CML cell lines), E-MTAB-16053 (RNA-seq of patients with CML in blast phase and chronic phase), E-MTAB-16054 (RNA-seq of K562 cells with MEF2C, MYB, MEIS1, and ZEB2 knockdown). The public data analysed in this project include GSM733656 (H3K27ac_Chip-Seq of K562), GSM733680 (H3K4me3_Chip-Seq of K562), GSM733692 (H3K4me1_Chip-Seq of K562); GSE4170 (Microarray data of BP-CMLs and CP-CMLs); EGAS00001003071 (the RNA-Seq data of BP-CMLs and CP-CMLs).

## Disclosure of Conflicts of Interest

None declared.

## Notes

### Competing Interest Statement

The authors have declared no competing interest.

