## Supplementary Materials for "The transcriptional regulatory circuit as a driver and therapeutic target in CML blast crisis"

#### Supplementary Methods

##### Antibodies and Reagents

The following antibodies were used in this study: Anti-MEIS1 (Abcam, ab19867), Anti-MEF2C (Proteintech, 10056-I-AP), Anti-ZEB2 (Proteintech, 67514-1-Ig), Anti-MYB (Abclonal, A13776), Anti-GAPDH (RM2002), Anti-AKT (Proteintech, P.10176), Anti-p-AKT (Proteintech,P.66444), Anti-GLUL (HUABIO, EM1902-39), Anti-VAV1 (HUABIO, EM1709-97), Anti-DOCK2 (HUABIO, HA721993), Anti-RhoA/B/C (SAB, #48641), Anti-p-STAT5 (Milipore， 2712453)， Anti-STAT5 (CST, 9363S), Anti-β-ACTIN (Proteintech, 205365-i-ap), Anti-BCR-ABL (CST, 2862S), Anti-p-BCR-ABL (Abcam, ab307142), Anti-Cleaved-Caspase3 (Abcam, AB2302), Anti-PARP (CST, 9542S), APC anti-human CD71(BD Pharmingen, 551374), APC anti-human CD61 (Proteintech, 65185), PE-Cy7 anti-mouse/human CD11b (BioLegend, 101216).

Reagents and kits included: Mebendazole (Selleck, s4610), D-Luciferin, Potassium Salt (Yeasen, 40902ES03), BioT (Bioland Scientific LLC, B01-00), Dual Luciferase Reporter Gene Assay Kit (Yeasen, 11402ES60), Hyperactive Universal CUT&Tag Assay Kit for Illumina (Vazyme, TD903-02), Cell Proliferation and Toxicity Test Kit (CCK-8) (Meilunbio, MA0218-L), Giemsa assay kit (Solarbio, G1010).

##### Chromatin Immunoprecipitation (ChIP)

$5\times10^6$ cells were collected and fixed with 1% paraformaldehyde for 10 minutes at room temperature. The fixation was terminated by adding an equal volume of 250 mM glycine. Samples were washed with PBS, then lysed with lysis/wash buffer on ice for 30 minutes. After centrifuge, cell pellets were harvested, resuspended in shearing buffer, and sonicated to shear genomic DNA to 300–500 bp. For immunoprecipitation, solubilized chromatin was rotated with target antibodies or IgG control antibody overnight at 4°C. Next, the antibody–chromatin complexes were pulled down by magnetic IgG beads, and the bound DNA was eluted with elution buffer. After decrosslink and RNA degradation, immunoprecipitated DNA was extracted using the Min-Elute PCR Purification Kit (Qiagen), followed by either qPCR analysis or DNA library preparation and sequencing on the HiSeq 4000 platform (Illumina).

##### ChIP-seq and CUT&Tag analysis

Initially, raw reads were aligned to the hg19 reference genome using Bwa (version 1.2.2) with default parameters[1]. Following this, high-quality mapped reads were filtered and sorted with SAMtools (version 1.3.1) using the “-q 10” option[2]. PCR duplicates were removed using the sambamba markdup tool (version 1.136), and regions identified as blacklisted were filtered out with bedtools (version 2.27.1).

For histone ChIP-seq analysis, peaks were called using MACS2 (Model-Based Analysis of ChIP-Seq, version 2.1.2) with the following parameters: “-q 0.01 –extsize 146 -nomodel -B”[3]. Bigwig files were created using the bamCompare function from deepTools (version 3.1.3) with parameters “–operation subtract –normalizeUsing CPM –extendReads 146 –binSize 20”[4]. For CUT&tag data, peaks were identified with MACS2 using the “-broad -q 0.01” options. Bigwig files for this data were generated via the bamCoverage function in deepTools with default parameters.

**CUT&Tag**

$15 \times 10^4$ cells were harvested and lysed with NE buffer. Samples were then centrifuged and resuspended in wash buffer. Nuclei were bound to ConA magnetic beads. The ConA-bound nuclei were then suspended and incubated with specific primary antibodies overnight at 4°C. After incubation, the samples were processed according to the manufacturer’s instructions for the CUT&Tag kit (TD903, Vazyme).

**RNA-Seq analysis**

Raw sequencing reads were aligned to the GRCh37 reference genome using HISAT2 (version 2.2.0) and subsequently quantified with the htseq-count tool (version 0.11.3) under default parameters. The raw read counts were normalized using FPKM. Differential expression analysis was conducted using the DESeq2 package (version 1.44.0), applying a threshold of $p\text{-value} < 0.05$ and an absolute log2 fold-change greater than 0.3.

**Plasmid construction and Lentivirus packaging**

The MEF2C, MEIS1, MYB, and ZEB2 expression vectors were amplified based on the pCDH-CMV-MCS-EF1-puro vector, and a 3xFLAG-tag was added via PCR. pLKO.1-puro vector expressing shRNAs targeting MEF2C, MEIS1, MYB, ZEB2, and GLUL were constructed and confirmed by DNA sequencing. The single guide RNAs (sgRNAs), targeting the enhancer site of ZEB2 , were inserted into lentiviral plasmid encoding dCas9 (pLV hU6-sgRNA hUbC-dCas9-KRAB-T2A-Puro, Addgene, #71236). The process involved designing sgRNA for MEF2C, MYB, MEIS1, and ZEB2 using CHOPCHOP, followed by directional cloning into the LentiCRISPRv2 vector. Positive clones were verified via Sanger sequencing and selected for plasmid purification. To produce viral particles, the pSPAX2 and pMD2.G packaging constructs were co-transfected into HEK293T cells. The supernatants containing viral particles were collected and filtered through a 0.45 μm filter after transfection for 48 hours. Cells of interest were then infected with the virus under 10 ug/mL polybrene for 48 hours. The sequences of shRNA, sgRNA were provided in Supplementary Table 2.

**Cell Proliferation Assay**

The pre-treated cells, which had been transfected with the indicated shRNAs or overexpressed genes, were seeded into 96-well plates at a density of $2\times10^3$ cells per well. Subsequently, 10 μL of CCK-8 reagent was added to each well at day 1 to day 5. After incubation at 37°C for 2 hours, absorbance was measured at 450 nm using a microplate reader (Thermo).

**Cell Toxicity Assay**

Cells were seeded into 96-well plates at a density of $1\times10^4$ cells per well and treated with a concentration gradient of MBZ or YKL-05-009 for 72 hours. Subsequently, 10 μL of CCK-8 reagent was added to each well, and after 2 hours of incubation, absorbance was measured at 450 nm using a 96-well microplate reader.

**RT-qPCR**

Total RNA was extracted using QIAGEN RNA extraction reagents (Cat. nos. 1018013, 79216, 1053394). For each sample, 1 μg of purified RNA was reverse-transcribed into complementary DNA (cDNA) using the Strand cDNA Synthesis SuperMix (Yeasen). Quantitative real-time PCR (qPCR) was performed on the CFX384 Real-Time PCR Detection System (Bio-Rad) with gene-specific primers and Universal Blue qPCR SYBR Master Mix (Yeasen). Gene expression levels were normalized to GAPDH and quantified using the comparative Ct method. The sequences of RNA primers are provided in Supplementary Table 3.

**Giemsa Staining**

Cells were harvested and resuspended in PBS. Once the labeled slides were prepared, an appropriate volume of the suspension was aspirated for smearing. After the smear was air-dried, methanol-fixed smears were treated with Giemsa reagent (Solarbio, China) for 30 minutes at room temperature following the manufacturer's instructions, followed by rinsing with running water. Subsequently, the dried smears were imaged using a light microscope (Leica Microsystems, Wetzlar, Germany).

**Dual Luciferase Reporter Assay**

Experimental procedures were adapted from previous studies[5]. Candidate enhancer or promoter regions were amplified by PCR and subsequently cloned into either the pGL3-Promoter luciferase reporter vector or the pGL3-Basic luciferase reporter vector (IGE). K562 cells were collected and resuspended in 100 μL of serum-free medium containing 10 μg of pGL3 and pRL-TK (4:1 ratio), then transfected by electroporation. pRL-TK vector was co-transfected as a normalization control. Forty-eight hours after transfection, luciferase activity was measured using the Dual-Luciferase Reporter Assay System (YEASEN).

**Cell Cycle Assay**

A certain number of cells were harvested, washed twice with PBS, and resuspended in pre-chilled 70%

ethanol, followed by incubation overnight at 4°C. Cell pellets were washed twice with PBS and stained with propidium iodide (PI) solution. Flow cytometry was performed to detect cell cycle distribution, and the data were analyzed using FlowJo X software.

**ELISA Assay of Glutamine**

$1\times10^6$ pre-treated cells were harvested, washed twice with PBS, and resuspended in 100 μL PBS. The samples were then processed according to the instructions provided with the glutamine kit (MB-3847A, MEIBIAO BIOLOGY). The abundance of glutamine was determined by the standard curve.

**Immunoblotting Assay**

Whole cell lysates were prepared in RIPA lysis buffer. After quantification, proteins were separated by SDS-PAGE and transferred to PVDF membrane. Immunoblotting was performed using indicated specific primary antibodies and horseradish peroxidase (HRP)-conjugated secondary antibodies.

**Co-Immunoprecipitation (Co-IP) Assay**

A certain number of cells were harvested and lysed on ice for 30 minutes. The supernatant was collected and divided into three portions: one portion served as input, while the other two were incubated overnight at 4°C with protein A/G magnetic beads conjugated to antibodies (IgG and the antibody against the target protein). The protein-beads complex were then washed three times with lysis buffer, and the bound proteins were eluted to proceed with immunoblotting.

**Gene Set Enrichment Analysis**

For the knockdown of MEIS1, MYB, ZEB2 and MEF2C, the down-regulated genes were individually chosen as the library genes. Then GSEAPreranked was performed using the fold change values from the knockdown of the respective TFs compared to the control as the ranked list. The significant terms with q value < 0.05 were shown.

**GST-pull Down Assay**

Lysates from CML cells were prepared in ice-cold lysis buffer (50 mM Tris-HCl pH 7.0, 150 mM NaCl, 10 mM $MgCl_2$, 1% Triton X-100, 10 mM Mg-acetate, protease inhibitor cocktail), centrifuged at 15,000 g for 20 min at 4°C. GTP-agarose beads (Sigma, #G9768) were washed three times with lysis buffer. Equal protein amounts of lysate were rotated with beads overnight at 4°C. Bead complexes were washed five times in lysis buffer (3,000 g, 3 min/wash), and bound proteins were eluted by boiling in 1× SDS sample buffer. GTP-bound (active) and total RhoA/B/C levels were analyzed by SDS-PAGE and immunoblotting using isoform-specific antibodies.

**Xenograft Assays of Blast CML in NSG Mice**

The experiments were subjected to the approval of the Institutional Animal Care and Use Committee of

Guangzhou Medical University and following the ethical rules of animal experiments (GY2021-089). Stable KBM5 cell lines engineered to express luciferase-GFP and shRNA constructs were intravenously injected into NSG mice via the tail vein ($5\times10^6$ cells/mouse)[6, 7]. Mice were divided into five experimental groups (n=15 per group): Scramble control, shMEF2C, shMYB, shMEIS1, and shZEB2. Body weight and bioluminescent tumor progression were monitored every other day using in vivo imaging. At the experimental endpoint, 5 mice/per group were euthanized, bone marrow mononuclear cells were isolated for flow cytometric differentiation analysis, cytospin preparations were generated, and protein lysates were prepared for Western blot analysis. Remaining mice were continuously monitored, and survival times were rigorously documented until all experimental endpoints were reached.

Separately, $5\times10^6$ luciferase-GFP-expressing KBM5 cells were intravenously inoculated into NSG mice (n=12). Upon detection of bioluminescent tumor signals, mice were randomized into two treatment groups (n=6 per group): vehicle (0.3% CMC-Na) and MBZ-treated (75 mg/kg/day via oral gavage). Tumor growth was monitored via bioluminescence imaging and body weight tracking. At study termination, peripheral blood leukocyte counts were determined, bone marrow mononuclear cells underwent flow cytometric analysis, cytospin slides were prepared, and protein lysates were collected for Western blot validation.

**Statistical Analysis**

Two-tailed Student's t-test was used for the statistics of differences in measurement data between groups, two-way analysis of variance (ANOVA) was used for statistics of differences in measurement data between multiple groups, and chi-square test was used for statistics of differences in count data between groups. All statistics and graphs were created with GraphPad Prism 8.0. Differences were considered statistically significant at $P < 0.05$ (*), $P < 0.01$ (**) , $P < 0.001$ (***) and $P<0.0001$ (****); ns, not significant.

5. Jiang L, Wang J, Peng G*, et al.* A Super-Enhancer-Driven Transcriptional Regulatory Circuit

Underlying Abiraterone Resistance in Castration-Resistant Prostate Cancer. *Adv Sci (Weinh)*, 2025; 12(31): e01284.

6. Carter B Z, Mak D H, Shi Y*, et al.* Regulation and targeting of Eg5, a mitotic motor protein in blast crisis CML: overcoming imatinib resistance. *Cell Cycle*, 2006; 5(19): 2223-9.
7. Valletta S, Dolatshad H, Bartenstein M*, et al.* ASXL1 mutation correction by CRISPR/Cas9 restores gene function in leukemia cells and increases survival in mouse xenografts. *Oncotarget*, 2015; 6(42): 44061-71.

### Supplementary Table

**Supplementary table 1**

| ID | Sex | Age | Diagnose | WBC | BM blast cells (%) | BCR-ABL1 | FISH |
|---|---|---|---|---|---|---|---|
| | | | | *10^9/L | | p210，p190，p230 | % postive |
| 1-CML-BP | FEMALE | 28 | CML-BP | 175.7 | 77.0% | P210 | N/A |
| 2-CML-CP | MALE | 44 | CML-CP | 110 | 1.0% | P210 | 91.7% |
| 3-CML-CP | MALE | 32 | CML-CP | 292.72 | 1.0% | P210 | 66.8% |
| 4-CML-CP | MALE | 24 | CML-CP | 45 | 1.5% | P210 | 90.0% |
| 5-CML-BP | MALE | 45 | CML-BP | 313.5 | 40% | p210 | N/A |
| 6-CML-CP | MALE | 41 | CML-CP | 142.68 | 1.0% | P190，P210 | 78.0% |
| 7-CML-BP | MALE | 63 | CML-BP | 105.6% | 70.0% | P210 | N/A |
| 8-CML-BP | FEMALE | 59 | CML-BP | 8.6 | 66.0% | P210 | N/A |
| 9-CML-BP | FEMALE | 73 | CML-AP | 0.99 | 21.0% | P210 | 38.5% |
| F804M_CML-CP | MALE | 53 | CML-CP | 44.08 | 3.0% | P210 | 100% |
| J6776M_CML-CP | FEMALE | 18 | CML-CP | 14.11 | 3.0% | P210 | 65.0% |
| L1459M_CML-CP | MALE | 27 | CML-CP | 125.8 | 0.5% | P210 | N/A |
| L3941M_CML-CP | MALE | 55 | CML-CP | 169.7 | 2.0% | P210 | N/A |
| L4344M_CML-CP | MALE | 40 | CML-CP | N/A | 1.0% | P210 | 35.0% |
| L5440M_CML-CP | MALE | 26 | CML-CP | 2.61 | 1.0% | P210 | N/A |
| L7831M_CML-CP | FEMALE | 35 | CML-CP | 269.5 | 2.0% | P210 | 85.0% |
| Y1074M_CML-CP | FEMALE | 26 | CML-CP | 19.12 | 4.0% | P210 | 95.0% |
| Y3939M_CML-CP | MALE | 36 | CML-CP | 70.67 | 3.0% | P210 | 99.0% |
| Y8720M_CML-CP | MALE | 40 | CML-CP | 5.27 | 2.0% | P210 | N/A |
| F4315m_CML-BP | FEMALE | 40 | CML-BP | 67.87 | 28.0% | P210 | 99% |
| F724M_CML-BP | FEMALE | 29 | CML-BP | 541.5 | 42.0% | P210 | 95% |
| J1848M_CML-BP | MALE | 33 | CML-BP | 17.36 | 35.0% | P210 | 30.0% |
| J7542M_CML-BP | FEMALE | 36 | CML-BP | 1.96 | 45.0% | P210 | N/A |
| J8918M_CML-BP | MALE | 26 | CML-BP | 3.48 | 55.5% | P210 | N/A |
| L9685M_CML-BP | MALE | 41 | CML-BP | 2.18 | 41.0% | P210 | 51.0% |
| Y283M_CML-BP | FEMALE | 52 | CML-BP | 658.19 | 41.0% | P210 | 100.0% |
| Y4299M_CML-BP | FEMALE | 59 | CML-BP | 53.68 | 39.0% | P210 | 100.0% |
| Y4946M_CML-BP | MALE | 70 | CML-BP | 99.49 | 37.0% | P210 | 99.0% |
| Y6033M_CML-BP | MALE | 51 | CML-BP | 212.97 | 41.0% | P210 | 98.0% |

**Supplementary table 2**

shRNA sequence of target genes

| Genes | | Sequence (5'-3') |
|---|---|---|
| Homo-ZEB2 | sh#1 | CCGGGACTGCAAGGCTGAAGAAATTCTCGAGAATTTCTTCAGCCTTGCAGTCTTTTT |
| | sh#2 | CCGGTCTGAAAGAACACCTGCGAATCTCGAGATTCGCAGGTGTTCTTTCAGATTTTT |

| | | |
|---|---|---|
| Homo-MEIS1 | sh#1 | CCGGCCAGCATCTAACACACCCTTACTCGAGTAAGGGTGTGTTAGATGCTGGTTTTT |
| | sh#2 | CCGGCGGTATATTAGCTGTTTGAAACTCGAGTTTCAAACAGCTAATATACCGTTTTT |
| Homo-MEF2C | sh#1 | CCGGCCCAATGAATTTAGGAATGAACTCGAGTTCATTCCTAAATTCATTGGGTTTTT |
| | sh#2 | CCGGGCCTAGAATTTGATACGCTTTCTCGAGAAAGCGTATCAAATTCTAGGCTTTTT |
| Homo-MYB | sh#1 | CCGGCCAGATTGTAAATGCTCATTTCTCGAGAAATGAGCATTTACAATCTGGTTTTT |
| | sh#2 | CCGGGCTATCAAGAACCACTGGAATCTCGAGATTCCAGTGGTTCTTGATAGCTTTTT |
| Homo-RUNX1 | sh#1 | CCGGCCTCGAAGACATCGGCAGAAACTCGAGTTTCTGCCGATGTCTTCGAGGTTTTTg |
| | sh#2 | CCGGGAACCAGGTTGCAAGATTTAACTCGAGTTAAATCTTGCAACCTGGTTCTTTTTg |
| Homo-SOX4 | sh#1 | CCGGGCGACAAGATCCCTTTCATTCCTCGAGGAATGAAAGGGATCTTGTCGCTTTTTGAATT |
| | sh#2 | CCGGGAAGAAGGTGAAGCGCGTCTACTCGAGTAGACGCGCTTCACCTTCTTCTTTTTGAATT |
| Homo-GLUL | sh#1 | CCGGGCACACCTGTAAACGGATAATCTCGAGATTATCCGTTTACAGGTGTGCTTTTTG |
| | sh#2 | CCGGCCAGGAGAAGAAGGGTTACTTCTCGAGAAGTAACCCTTCTTCTCCTGGTTTTTG |
| Homo-HOXA9 | sh#1 | CCGGCGCTGTACCCGCTGCGGTGTACTCGAGTACACCGCAGCGGGTACAGCGTTTTT |
| | sh#2 | CCGGCGCCTCGTGGAACCCAGTGCACTCGAGTGCACTGGGTTCCACGAGGCGTTTTT |

CRISPR-Cas9 sgRNA sequence of target genes

| **Genes** | **Sequence (5'-3')** |
|---|---|
| homo-MEF2C-KO | GTGTTGTGGGTATCTCGAAGGGG |
| homo-MYB-KO | CGTCGGAAGGTCGAACAGGAAGG |
| homo-MEIS1-KO | ACGGCATCTACTCGTTCAGGAGG |
| homo-ZEB2-KO | CAGAGCTTGGCTCACGTGTGGGG |

CRISPR-dCas9 sgRNA sequence of ZEB2-E2

| Genes | | Sequence (5'-3') |
|---|---|---|
| ZEB2-E2 | sg#1 | GTTGATATGGACGGTCTGCG |
| | sg#2 | AGGTTCCACGCAACTGTTGG |

**Supplementary table 3**

qPCR primers of target genes

| **Genes** | **Forward (5'- 3')** | **Reverse (5'- 3')** |
|---|---|---|
| Homo-VAV1 | AGCAGAAGGACTGTACCGGA | TGAAGGGGAACTGCAAGGTG |

| | | |
|---|---|---|
| Homo-DOCK2 | AGATGGGTGACCAGCACTAC | GCCATCCAGTCTCCAGGGTA |
| Homo-MYB | ATCTCCCGAATCGAACAGATGT | TGCTTGGCAATAACAGACCAAC |
| Homo-ZEB2 | GGAGACGAGTCCAGCTAGTGT | CCACTCCACCCTCCCTTATTTC |
| Homo-MEF2C | CCAACTTCGAGATGCCAGTCT | GTCGATGTGTTACACCAGGAG |
| Homo-MEIS1 | GCGCAAAGGTACGACGATCT | GGTACTGATGCGAGTGCAGA |
| Homo-HOXA9 | AAAAACAACCCAGCGAAGGC | ACCGCTTTTTCCGAGTGGAG |
| Homo-SOX4 | GACCTGAACCCCAGCTCAAA | AGCCGGGCTCGAAGTTAAAA |
| Homo-RUNX1 | CCTCAGGTTTGTCGGTCGAA | CTTGCGGTGGGTTTGTGAAG |
| Homo-GLUL | CTCTCGCGGCCTAGCTTTAC | ACTTGCTGAGGTGGTCATGG |

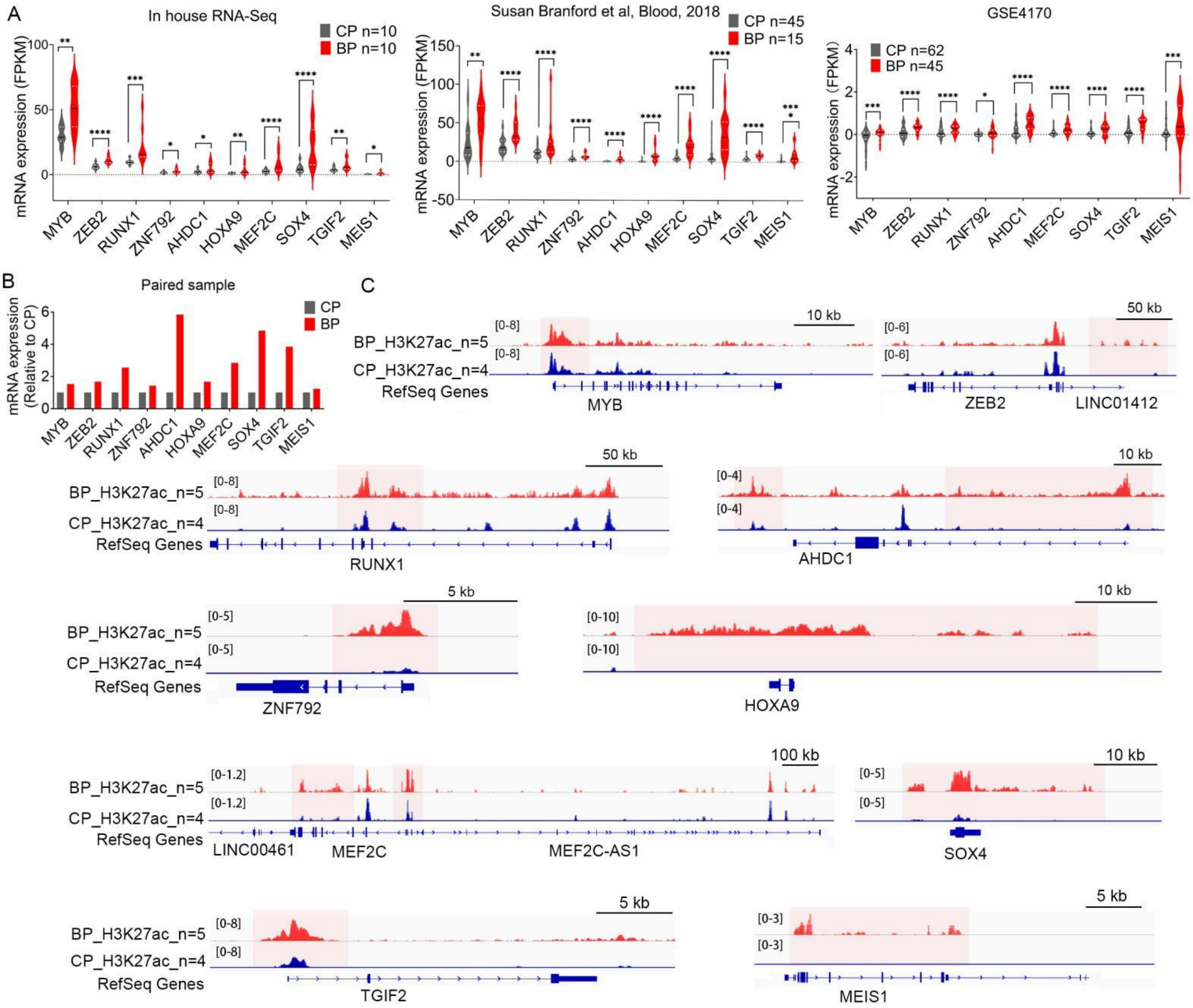


**Supplementary Figure 1**

A. The mRNA expression of candidate TFs in CP-CMLs and BP-CMLs.

B. The mRNA expression of candidate TFs in paired CP-CML and BP-CML.

C. The tracks show profiles of ChIP-Seq signals of H3K27ac at candidate TFs gene loci.

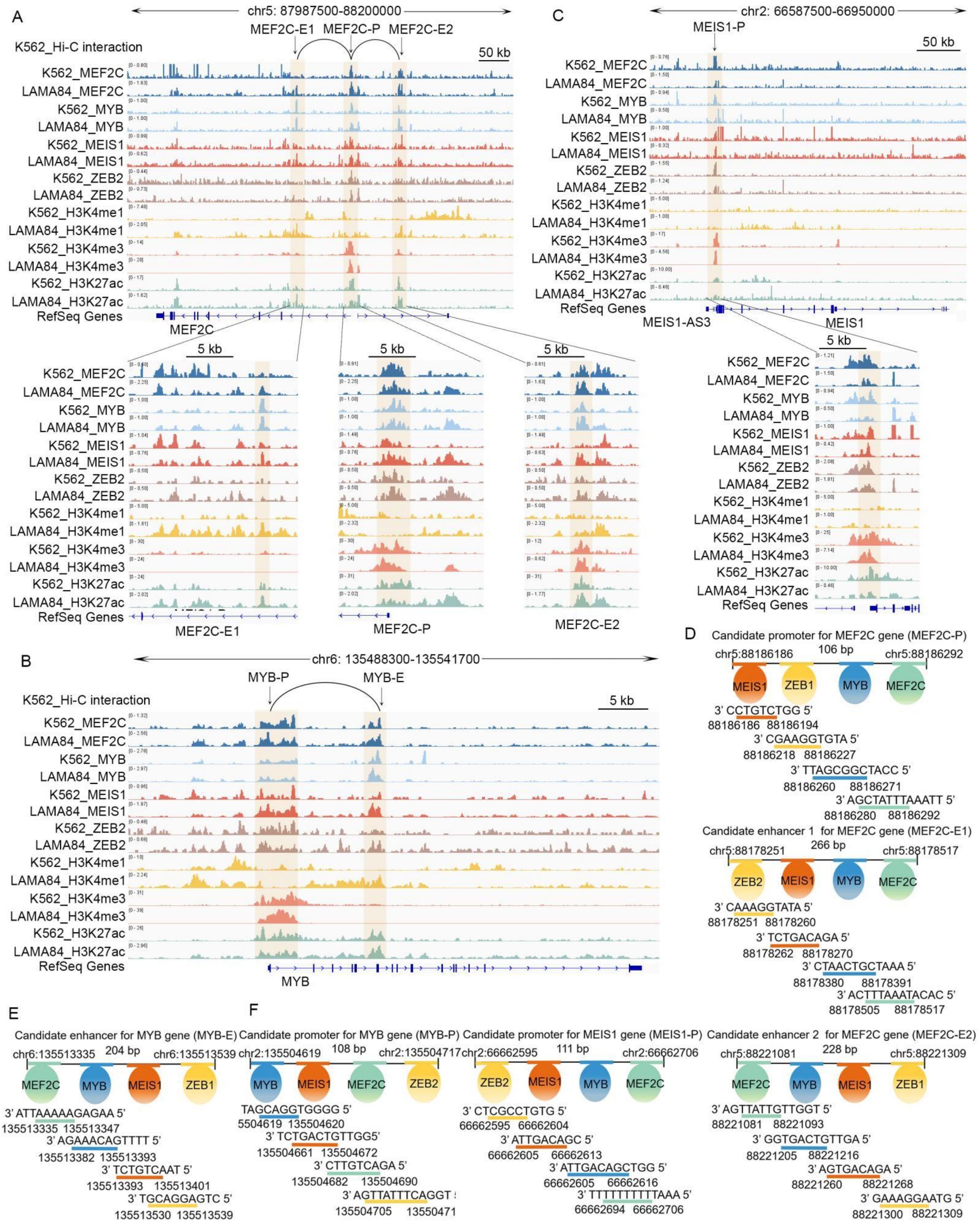


**Supplementary Figure 2**

The tracks show profiles of ChIP-Seq or CUT&Tag signals of indicated antibodies at MEF2C (A), MYB (B), and MEIS1 (C) gene loci.

The schematic representation of the nearest distance pattern of MEF2C (D), MYB (E), and MEIS1 (F) motifs at promoter and enhancer.

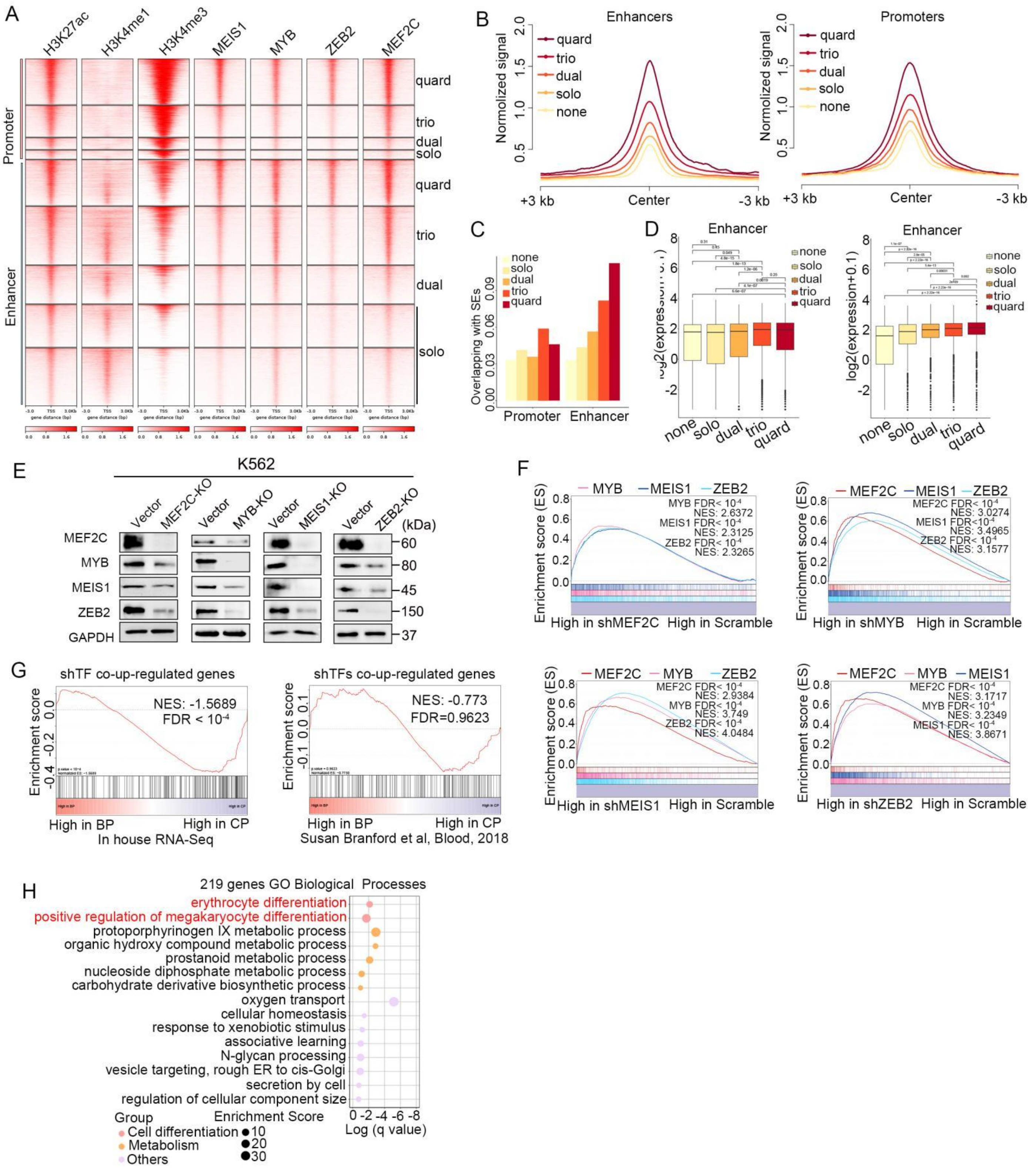


**Supplementary Figure 3**

A. Heatmaps of CUT&Tag signals of indicated factors in LAMA84 cells, stratified by different combinatorial binding patterns.

B. The normalized signal intensity of H3K27ac among the peaks of indicated groups.

C. The proportion of super enhancers (SEs) among peaks in the indicated groups.

D. The mRNA expression of adjacent to the indicated peaks in LAMA84 cells.

E. The protein level of TFs with CRISPR-CAS9 mediated knock out in K562 cells.

F. Genes up-regulated by knocking down each TF are subjected to GSEA enrichment analysis with the RNA-Seq data of the other three knockdown.

G. Genes co-up-regulated by knocking down MEF2C, MYB, MEIS1, ZEB2 are subjected to GSEA enrichment analysis with the RNA-Seq data of BP and CP-CMLs.

H. GO biological enrichment of genes co-up-regulated in MEF2C, MYB, MEIS1and ZEB2 knockdown.

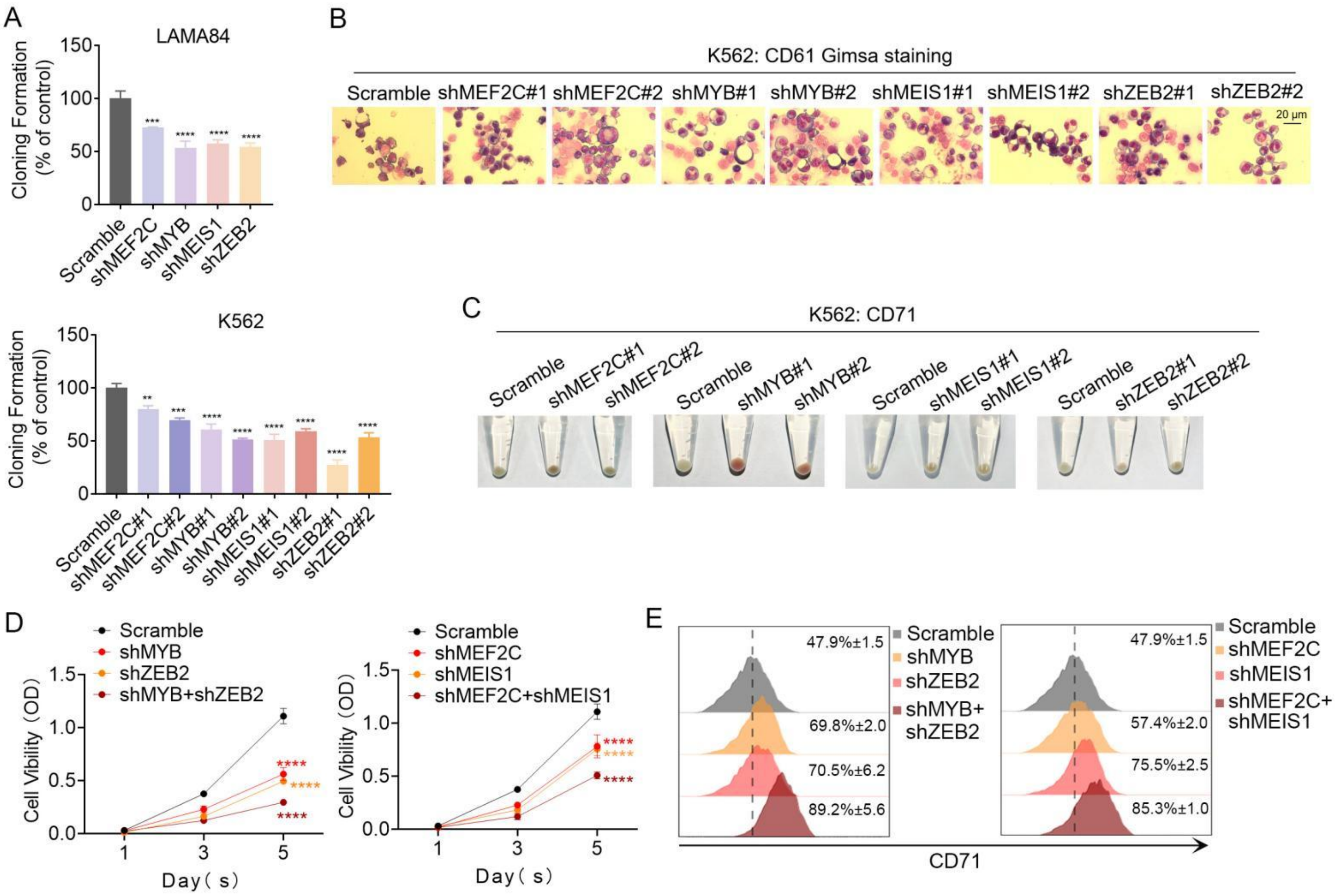


#### Supplementary Figure 4

A. The colony formation of MEF2C, MYB, MEIS1, ZEB2 knockdown and Srcamble at 14 days after plate.

B. The represent image of Gimsa staining after 1 nM PMA treatment 24h at K562 cells (n=3).

C. The represent image of Gimsa staining after 10 μM hemin treatment 48h at K562 cells (n=3).

D. Cell viability of knockdown one or two indicating TFs in K562.

E. Cell differentiation were detected by flow cytometer with one or two indicating TFs knockdown, CD71 antibody for erythroid differentiation (10 μM hemin treatment 48h).

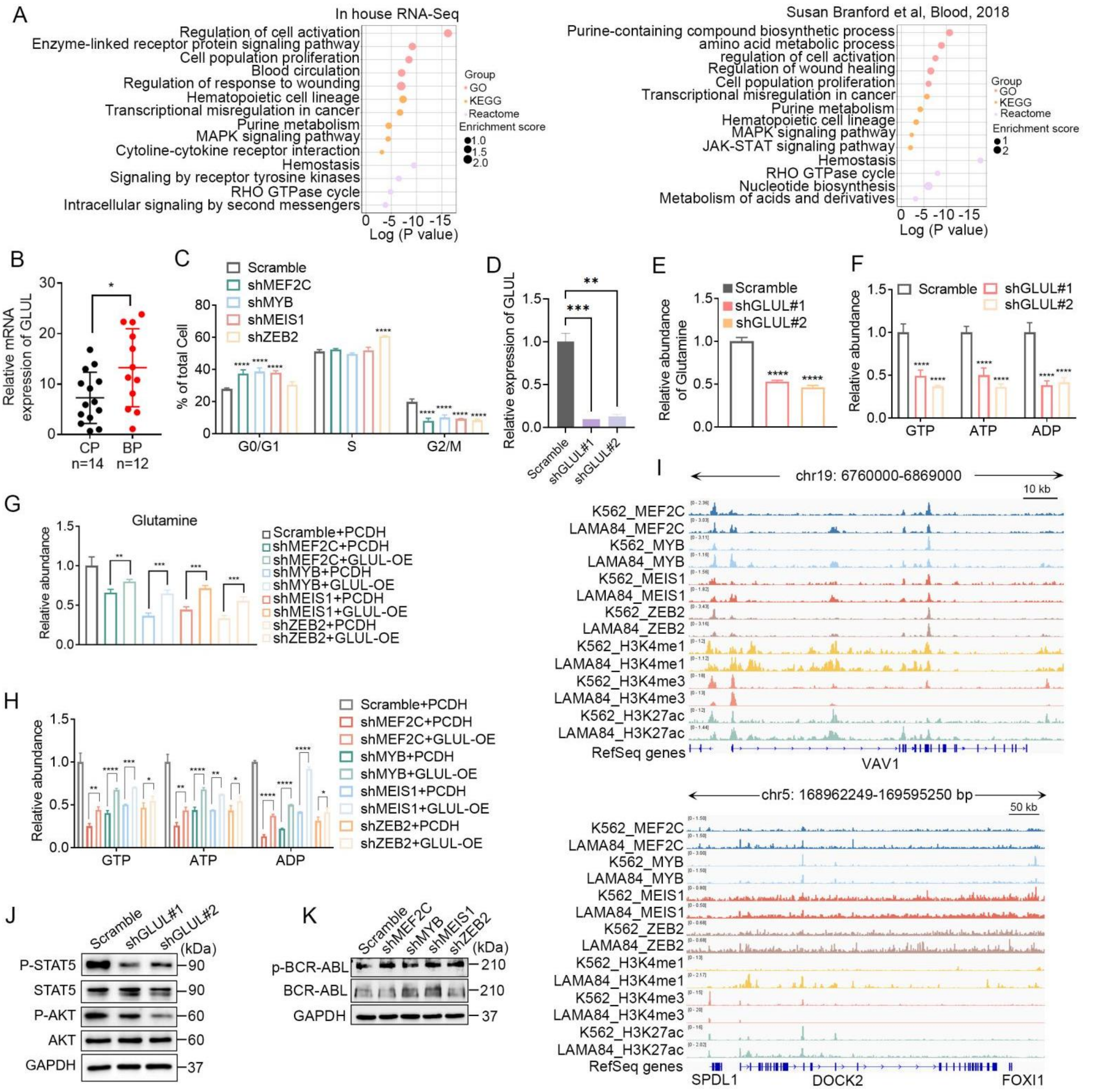


**Supplementary Figure 5**

A. The pathway enrichment of genes up-regulated in BP-CML.

B. mRNA expression of GLUL in CP-CML(n=14) or BP-CML(n=12).

C. The distribution of cell cycle under TFs knockdown.

D. mRNA expression of GLUL knockdown in K562 cells.

E. The abundance of Glutamine measured by ELISA assay with GLUL knockdown and Scramble.

F. The abundance of GTP, ATP, and ADP measured by HLPC with GLUL knockdown and Scramble.

G. The abundance of Glutamine evaluated in K562 cells with TF knockdown and Scramble, followed by lentiviral overexpression of PCDH or GLUL.

H. The abundance of GTP, ATP, and ADP evaluated in K562 cells with TF knockdown and Scramble, followed by lentiviral overexpression of PCDH or GLUL.

I. The tracks show profiles of CHIP-Seq or CUT&Tag signals of indicated antibodies at DOCK2 and VAV1 gene loci.

J. The protein level of indicated factors were detected under GLUL silencing.

K. The protein level of indicated factors were detected under TFs silencing.

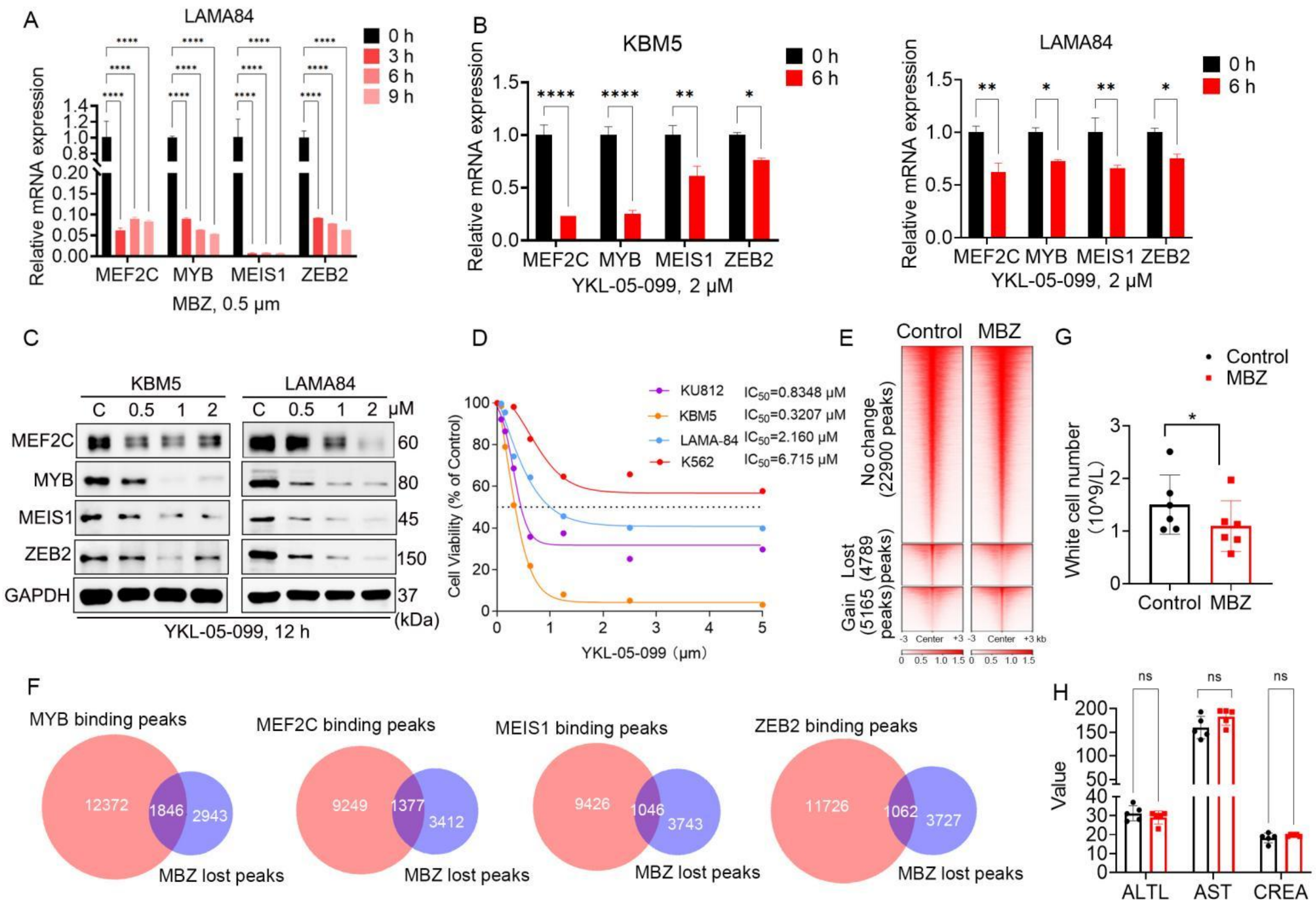


**Supplementary Figure 6**

A. LAMA84 cells were treated with Mebendazole, and then the indicated mRNA were detected by RT-qPCR.

B. KBM5 and LAMA84 cells were treated with YKL-05-099, and then the indicated mRNA were detected by RT-qPCR.

C. KBM5 and LAMA84 cells were treated with YKL-05-099, and then the indicated proteins were detected by Western blots.

D. Treatment of CML cells with YKL-05-099, MTS assay performed to measure cell viability after 72 h.

E. Heatmaps showing changes of ChIP-seq H3K27ac signals upon given of Mebendazole in K562 cells.

F. The overlap of TF binding peaks and the lost H3K27ac peaks upon administration of Mebendazole.

G. Total number of white cells in the Peripheral Blood of NSG mice with Mebendazole treatment (n=6).

H. The serum ALT, AST and CR levels were measured in Mebendazole-treated group (n=6). ALT: Alanine Aminotransferase; AST: Aspartate Aminotransferase; CR: Creatinine.
